# A gene-edited chicken model of ocular coloboma reveals Netrin-1 is required for pioneer cell detachment during optic fissure closure

**DOI:** 10.64898/2026.11.25.768265

**Authors:** Teresa Requena Navarro, Holly Hardy, Hannah Perry, Taran Kaur, Brian Ho Ching Chan, Jason Ioannidis, Dominique Meunier, Alix B E Johnston, Anna Raper, Mike J McGrew, Donald R Dunbar, Rachel Queen, Joe Rainger

## Abstract

Coloboma, a leading cause of childhood blindness, results from failed OFC. Variants in NTN1 (Netrin-1) are linked to coloboma and other tissue fusion defects, but how Netrin-1 influences epithelial fusion is unknown. We generated a germline NTN1 knockout chicken and analysed the fissure margins by immunofluorescence, in situ hybridisation, single-nucleus RNA sequencing and micro-CT. NTN1^−/−^ embryos developed highly penetrant bilateral colobomas. The basement membrane (BM) was remodelled normally, yet pioneer cells at the optic fissure margins (OFMs) failed to detach and mix across the midline, retaining apical actin and displaying increased cadherin at the fissure edges. The ventral neural retina showed reduced apoptosis and became overgrown and folded at the fissure edges. These data show that Netrin-1 is required for pioneer cell detachment and this process is independent of BM remodelling. We found that pioneer cells in wild type (WT) OFMs were transiently exposed to fibronectin-rich periocular mesenchyme (POM) during BM remodelling, with fibronectin and laminin fragments localised at their exposed edges. Netrin-1 reduced adhesion of HEK293 cells to fibronectin in vitro, with and without laminin, but not to laminin alone, suggesting Netrin-1 mediates a substrate specific effect on cell adhesion. ITGB8 and ITGAV were co-expressed with NTN1 in pioneer cells, and Netrin-1 co-immunoprecipitated with ITGB8, identifying integrin αVβ8 as a candidate Netrin-1 receptor in OFC. Integrin αVβ8 with fibronectin typically activate canonical TGFβ signalling, however we found pSMAD2 was specifically absent from pioneer cell nuclei during fusion. In contrast, phosphorylated FAK was present in WT but not mutant fissures, indicating that Netrin-1 activates integrin mediated focal adhesion signalling in OFC. We propose that Netrin-1 and αVβ8 enable pioneer cells to respond to transient POM-derived cues, such as fibronectin, triggering detachment and initiating fusion. This proposed mechanism could explain a direct role for the POM in mediating OFC.

## INTRODUCTION

Coloboma is a congenital malformation of the eye caused by failure of OFC and is a leading cause of defective vision in childhood (Chang et al., 2006; Lingam et al., 2021; Onwochei et al., 2000). In normal stages of early development, the ventral optic cup and stalk form a structural gap referred to as the optic (or choroid) fissure along their entire proximal-distal (PD) axis. As the eye grows, OFMs come together and fuse bidirectionally to complete the 3D circumference of the eye (Chan et al, 2021). Failure of this fusion results in colobomas at any region along the PD axis, resulting in persistent gaps at the iris, choroid or optic nerve, or a combination of all these (Chang et al., 2006; Lingam et al., 2021). Coloboma is genetically heterogeneous, with over 40 genes, and several environmental factors also linked to causality (ALSomiry et al., 2019; Gregory-Evans et al., 2004; Patel and Sowden, 2019; Selzer et al., 2021). However, most cases of coloboma (∼70-80%) remain genetically unresolved (Harding et al., 2022; Morrison et al., 2002; Owen et al., 2022; Williamson and FitzPatrick, 2014), which reflects how little is known about the molecular pathways and cell behaviours required at each step of OFC.

Developmental epithelial fusion events shape tissues such as neural tube, palate, inner ear and ventral body wall, and their failure underlies common congenital disorders (Ray and Niswander, 2012). In OFC, the opposing retinal epithelia edges first meet at their basal surfaces, which are preliminarily enclosed by a laminin rich BM (Gestri et al., 2018; Hardy et al., 2019). Fusion then requires several steps: breakdown or displacement of the BM at the epithelial edges, detachment of epithelial “pioneer cells” from their neighbours at these marginal edges (Eckert et al., 2020; Hardy et al., 2019) and their mixing across the midline, re-establishment of epithelial polarity and cell junctions in the fused seam, and BM remodelling to bond the fused epithelia (Chan, 2021). The inter-relatedness and dependence of each of these steps, and how these are orchestrated by specific genes and pathways, remains largely unknown. However, transcriptional profiling of optic fissure cells in human, mouse, zebrafish and chicken eyes has identified multiple coloboma candidate genes (Brown et al., 2009; Chan et al., 2025; Hardy et al., 2019; Patel et al., 2020; Richardson et al., 2019; Trejo-Reveles et al., 2023). These studies have consistently implicated epithelial to mesenchymal transition (EMT) -like pathways are active during fusion, but none have defined the function of these genes precisely within the fusion process.

The fusion candidate NTN1 encodes Netrin-1, a secreted laminin-related protein best known as an axon guidance cue (Lai Wing Sun et al., 2011). Netrin-1 was first implicated in tissue fusion in Ntn1^-/-^ mice with abnormally fused semicircular canal epithelia (Salminen et al., 2000). NTN1 is also highly expressed in the OFMs of zebrafish, chicken, human and mouse pioneer cells during OFC (Hardy et al., 2019; Richardson et al., 2019, Patel et al., 2020). In each species its expression is dynamic: enriched in pioneer cells before fusion then rapidly downregulated once fusion is complete. Genetic loss of function for Netrin-1 causes colobomas in mouse and zebrafish (Hardy et al., 2019; Richardson et al., 2019), with eye size and OFM apposition in mouse mutants appearing normal, suggesting defective fusion rather than growth is the underlying aetiology of the colobomas. These mice also feature cleft palates, collectively supporting a conserved requirement for Netrin-1 in fusion for multiple tissues. A coding variant in NTN1 has since been reported in a single patient with coloboma (Toms et al., 2025), suggesting this function extends to human development. Netrin-1 is enriched in extracellular matrix (ECM) and BMs and functionally interacts with cell receptors including DCC, Neogenin, DSCAM and integrins (Chaturvedi and Murray, 2021; Cirulli and Yebra, 2007; Lai Wing Sun et al., 2011; Yebra et al., 2003). However, Netrin-1 binding partners during fusion, and how these mediate the necessary downstream cellular mechanisms, have not been determined.

Here we developmentally model coloboma by generating a stable gene-edited NTN1 knockout chicken line using emerging avian genetic tools (Davey et al., 2018; McGrew et al., 2025). Homozygous embryos lacking Netrin-1 developed highly penetrant bilateral colobomas. During OFC their BM was remodelled normally, yet pioneer cells failed to detach and mix across the midline, which revealed that BM removal is not sufficient for fusion. We found that normal BM breakdown transiently exposed pioneer cells to the adjacent POM, which expresses fibronectin, and that Netrin-1 reduces cell adhesion to fibronectin in vitro. NTN1 was co-expressed with integrin αVβ8 in the OFM, and we found that Netrin-1 interacted with ITGB8 at the protein level. Analysis of integrin signalling found that pioneer cells are typically absent for nuclear pSMAD2, but that NTN1 mutant pioneer cells lacked the focal adhesion signalling seen in control tissues. We propose that Netrin-1, integrin αVβ8, and POM-derived factors such as fibronectin may act together to initiate pioneer cell detachment, uncovering a direct role for the POM in mediating OFC.

## RESULTS

### Generation of NTN1 knockout chicken line

We applied CRISPR/Cas9 to target the chicken NTN1 gene locus (NCBI Gallus gallus; ID:396389) in chicken primordial germ cells (PGCs)(Fig.1A), using a sgRNA we previously used to generate a knock-in GFP-reporter allele of NTN1 (Chan et al., 2025). We generated clonal male PGC populations and selected a clone with a homozygous 1-bp deletion that predicted a coding frameshift in exon 1 (p.Arg24fsX6), and nonsense-mediated decay of NTN1 mRNA (Fig.1B). PGCs for this allele (NTN1^-/-^) were injected into iCaspase9 (DAZL^iCaspase9^) surrogate host embryos (Ballantyne et al., 2021) to generate generation 0 (G0) founder chickens as previously described (Chan et al., 2025). Males were grown to sexual maturity and crossed to WT hens, and these G1 eggs were then incubated to obtain embryos at various developmental stages, or to hatch live birds for further breeding. We determined inheritance of the targeted allele and the viability and fertility of the line; all progenies of the G0 male were heterozygous (NTN1^+/-^) (n>175), with high egg fertility (>96.5%) and high viability of embryos (>93%) (Suppl.Fig.S1). We collected optic fissure and neural tube tissues from NTN1^+/-^ embryos and confirmed 50% reduction for both NTN1 mRNA and Netrin-1 protein compared to stage-matched wild type (WT) embryos (Suppl.Fig.S1). We then hatched and raised male and female NTN1^+/-^ chickens to sexual maturity, which displayed no observable physical or behavioural phenotypes. In G1 x G1 crosses to obtain homozygous offspring, we observed grossly normal development in WT, NTN1^+/-^, and NTN1^-/-^ embryos at Hamburger Hamilton (HH) stage HH28 (∼ embryonic day 6); Suppl.Fig.S1) with all three genotypes observed at expected mendelian rates (Fig.1C). Netrin-1 protein was undetectable in NTN1^-/-^ embryos compared to WT using western-blotting from whole-embryo samples (Fig.1D), and by immunofluorescence on cryosections from the optic fissure and neural tube floorplate (Fig.1E-F). Thus, we concluded that the NTN1^-/-^ allele (p.Arg24fsX6) is a complete loss of function (LOF) or “knockout” for Netrin-1. Similarly, we crossed our previously generated NTN1^T2A-GFP/+^ birds (Chan et al., 2025) to generate homozygote embryos and also confirmed loss of detectable Netrin-1 protein in NTN1^T2A-GFP/T2A-GFP^ embryos and observed the expected mendelian distributions of all three genotypes (NTN1^T2A-GFP/T2A-GFP^, NTN1^T2A-GFP/+^, and WT) (Suppl.Fig.S1).

**Fig. 1.**
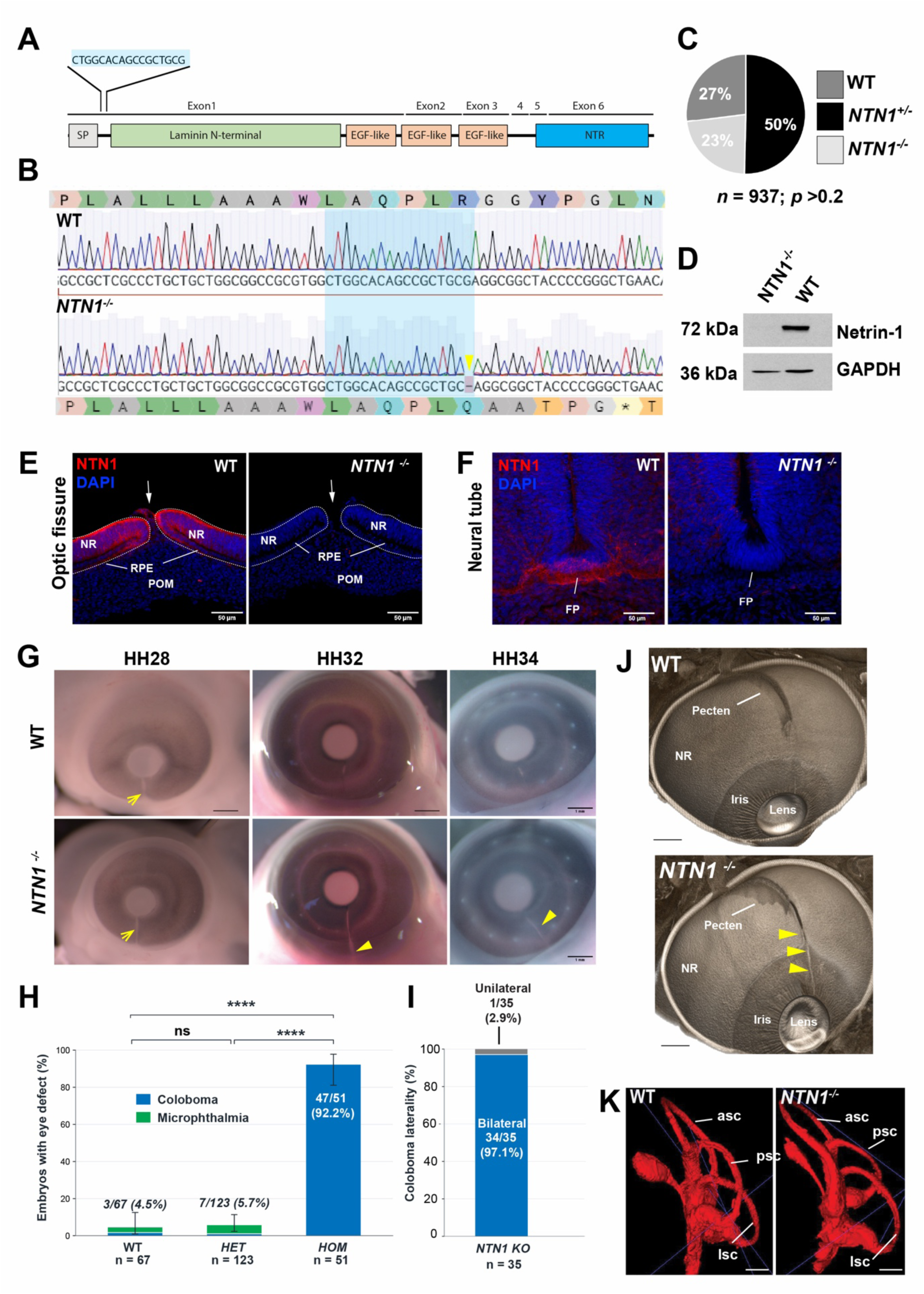
Generation of gene edited NTN1^-/-^ chicken embryos with highly penetrant ocular colobomas. (A) Gene editing strategy using sgRNA targeted to the signal peptide (SP) encoding region of NTN1 exon 1. (B) Aligned DNA chromatograms at the targeted locus for WT and NTN1^-/-^ PGCs, confirming 1bp deletion (arrow). The deletion results in a frame shift at the amino acid level and predicts a prematurely truncated Netrin-1 protein (p.Arg24fs6X). (C) Distribution of genotypes identified in NTN1^-/+^ X NTN1^-/+^ crosses. No statistical difference between expected and observed embryos. (D) Western blotting was unable to detect Netrin-1 protein in NTN1^-/-^ embryonic tissues. (E) Immunofluorescence for Netrin-1 in OFMs (HH28/E6) and (F) cranial neural tube (HH25/E5) cryosections confirmed loss of detectable Netrin-1 in NTN1^-/-^ embryos. (G) Whole eyes showing persistent colobomas (arrowheads) in NTN1^-/-^ embryos. Open arrows indicate naturally open OFM prior to fusion in both genotypes. (H) Quantitation of eye defects in WT (n=67; 1 unilateral coloboma), heterozygous (HET; n=123; 1 unilateral coloboma) and homozygous (HOM; n=51) at stages >HH34/E9 (**** p< 0.0001). (I) Laterality of coloboma in NTN1 KO embryos (n=35 embryos with coloboma). Error bars = 95% Clopper-Pearson CIs. (J) Soft tissue µCT eye scanning identified coloboma (arrowheads) extended from iris (anterior) to pecten (posterior) in NTN1^-/-^ embryos. (K) µCT scan reconstructions of semicircular canals identified no gross anatomical differences between WT and NTN1^-/-^. Abbreviations: NR, neural retina; RPE, retinal pigmented epithelium; POM, periocular mesenchyme; FP, floor plate of neural tube; asc, psc, lsc, are anterior-, posterior-, and lateral-semicircular canals, respectively.

### NTN1 knockout chickens exhibit highly penetrant ocular colobomas

Given both mouse and zebrafish knockout lines for Netrin-1 exhibit colobomas (Chan et al., 2025; Richardson et al., 2019) we assessed our NTN1 homozygous chicken embryos (NTN1^T2A-GFP/T2A-GFP^ and NTN1^-/-^) for this phenotype. The chicken OFM is open at HH28 but is fully fused by HH34 (E9) (Hardy et al., 2019). OFM anatomy was equivalent at HH28, and gross embryonic anatomy appeared normal in all genotypes (Suppl.Fig.S1). However, a persistent gap in the ventral retina was observed at all later stages (HH32 onwards) in NTN1^-/-^ eyes (Fig.1G; Suppl.Fig.S1). The colobomas in NTN1^-/-^ embryos were observed with penetrance >92% (47/51; Fig.1H) and were bilateral in >97% of embryos with colobomas (34/35; Fig.1I). Colobomas with similar phenotype anatomy, penetrance, and laterality were also observed for homozygous NTN1^T2AeGFP/2AeGFP^ embryos (Suppl.Fig.S1), indicating that both lines represent LOF/knockout alleles for NTN1. Homozygote birds with colobomas from both lines were then tested for eye size and were found to not show any significant difference compared to controls (i.e. not microphthalmic) (Suppl.Fig.S1). Whole embryo phenotyping using micro computed tomography (µCT) allowed non-destructive analyses for further fusion defects in NTN1^-/-^ embryos, first revealing the colobomas extended along the entire proximal-distal axis of the ventral eye, from the iris to the pecten (n=3; Fig.1J). µCT and 3D reconstruction analysis of the inner ear showed no gross fusion defects in NTN1^-/-^, with the gross anatomy of all three semicircular canals appearing normal and comparable to WT (n=3 embryos; 6 ears per genotype; Fig.1K). Optic chiasms and optic nerves also appeared grossly normal (Suppl.Fig.S1), and we also observed no changes to the palates of NTN1^-/-^ embryos (data not shown), however this was not surprising as the chicken secondary palate does not fuse during normal development. Thus, loss of Netrin-1 causes highly penetrant non-microphthalmic colobomas in chicken embryos, illustrating the evolutionarily conserved and essential requirement for NTN1 during OFC. This line is, to our knowledge, the first germline genetically engineered chicken with a developmental ocular defect.

### NTN1^-/-^ embryos exhibit failure of pioneer cell detachment and neural retina overgrowth

During active fusion stages (HH30), we observed that in NTN1^-/-^embryos BM remodelling occurred comparably with that observed in WT OFMs (Fig.2A). However, this BM modulation occurred in the absence of pioneer cell reorganisation and cell mixing across the opposing epithelia, with the folding points at the apical aspects of pioneer cells remaining apparently unmoved. Prior to fusion of the opposed OFMs, the laminin-rich BMs are organised juxtaposed to the basal cell edges of pioneer cells, whereas the pioneer cell’s apical regions are indicated by an enrichment of F-actin. To confirm the failure of pioneer cell reorganisation we stained for F-actin and found persistent apical enrichment in NTN1^-/-^ OFMs at their folding points compared to WT (Fig.2B), where F-actin was initially enriched at the folding point then became apically continuous in the fused retina. These data indicated that pioneer cells failed to detach from the folding point, despite BM remodelling. Staining for neurofilament (NF145) indicated that fusion failure was not due to ingression of ectopic ganglion cell axonal expansion into the pioneer domain (Suppl.Fig.2). We then analysed the consequence of this phenotype in later stage fissures. In WT fissures at HH34 (2 days post-fusion) the ventral retina was completely fused into continuous retinal pigmented epithelium (RPE) and neural retina (NR) epithelia at the midline (Fig.2C). In contrast, NTN1^-/-^ fissures appeared folded and lacked epithelial continuity at the midline. A single continuous layer of F-actin was observed at both the ganglion cell layer (GCL) and the apical-most region of the retina (the outer nuclear layer) facing the RPE in WT retinas, and the inner layer was covered by a laminin-rich BM (Fig.2D), and nuclei were organised into RPE and NR epithelial layers. In contrast, NTN1^-/-^ eyes at the same stage were defined by a continuous laminin-rich BM overlying the inner layer of the fissure, and there was F-actin enrichment observed at the GCL region comparable to WT, but cells at the fissure edges also exhibited persistent F-actin enrichment at their apical regions (Fig.2D) and the folding points appeared to be intact. At HH40 (3 days post-fusion) the WT retina was continuous at the midline and had begun clear retinal lamination, with multiple cell layers visible by light microscopy (Fig.2E). However, at the same stage the NTN1^-/-^ OFM appeared to still not have fused across the midline and was characterised by tissue overgrowth and folding. Of note, the RPE and NR on each side of the NTN1^-/-^ midline appeared as a continuum (Fig.2E; Suppl.Fig.S2), indicating that the pioneer cells in this region had not detached or progressed fusion. The folded tissue appeared to be a continuation of the neural retina rather than RPE as no pigment was observed, and the cells appeared to show organisation consistent with retinal lamination. This was confirmed by fluorescence in situ hybridisation for the inner nuclear layer marker OTX2 and the ventral outer nuclear layer marker VAX1 (Fig.2F), and by immunofluorescence for the neural retina markers SOX2 and PAX6 (Fig.2G-H). Thus, in the absence of Netrin-1, fissure margin pioneer cells do not undergo cell detachment or mixing across the midline, despite the reorganisation of the overlying BM. This novel phenotype illustrates that these two requisites of fusion – (i) pioneer cell detachment and reorganisation and (ii) BM remodelling, are separate events and are not interdependent. This data also implies that the epithelial behaviours needed for OFC require active initiation via Netrin-1 mediated mechanisms, rather than being an automatic process after BM removal. Our data also show that Netrin-1 plays no essential role in BM modulation during fusion.

**Fig. 2.**
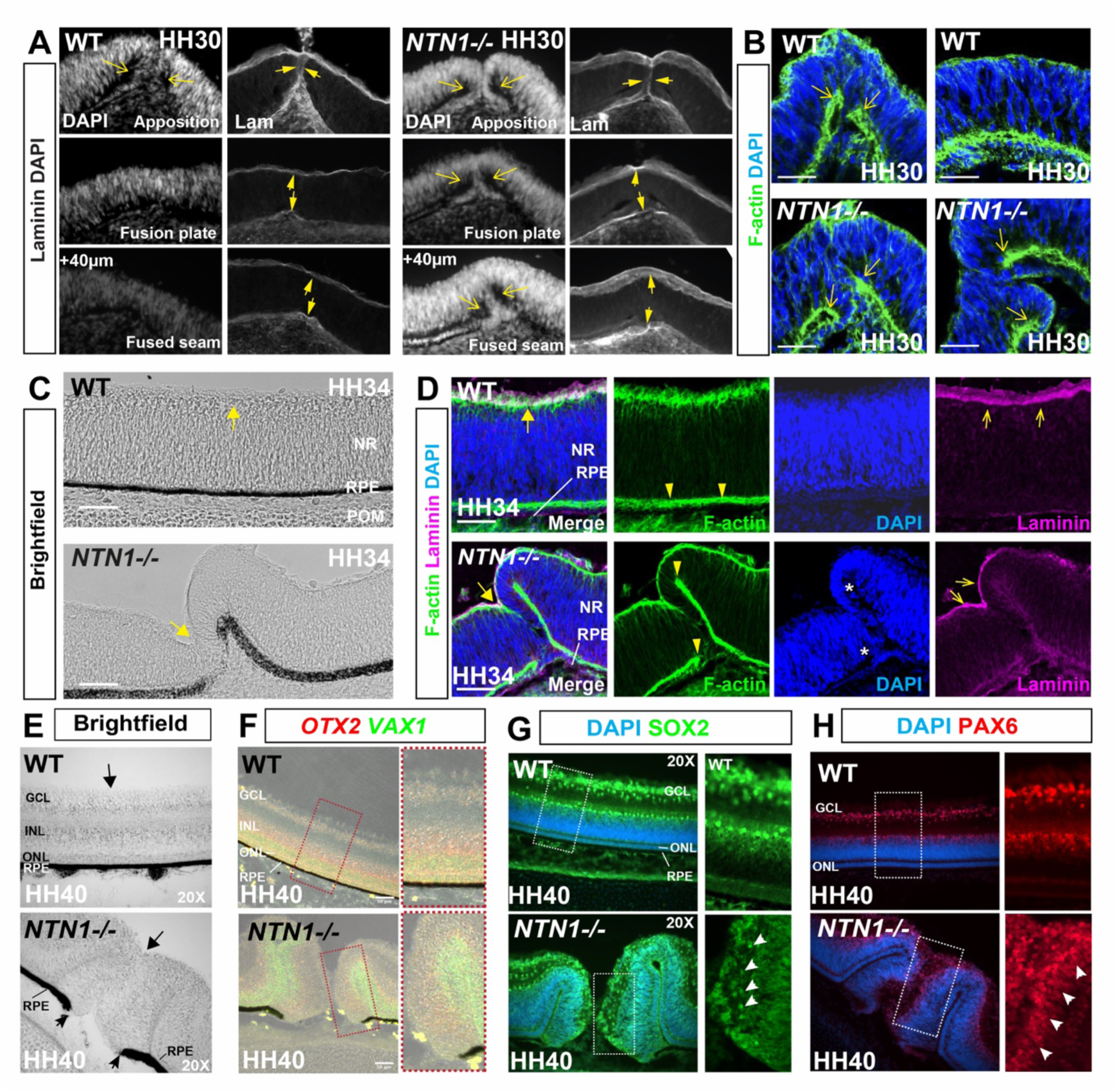
BM remodelling occurs but pioneer cells do not detach in NTN1^-/-^. (A) Nuclear staining (DAPI) and anti-Laminin immunofluorescence in WT illustrates BM (arrowheads) originally surrounds opposing basal edges of OFM epithelia but then is remodelled to be continuous with both the inner limiting membrane and the exterior face of the RPE, this coincident with pioneer cell mixing across the midline (folding point = open arrows). In NTN1^-/-^ the BM is remodelled (arrowheads), but pioneer cells do not mix across the midline, and their folding points are maintained in the fusion plate (open arrows). (B) F-actin staining illustrated continued maintenance of the folding point (arrows) in NTN1^-/-^ OFMs compared to WT. (C) Post-fusion OFMs show folding defect observed in NTN1^-/-^ compared to epithelial continuum in WT (Arrows = OFM midline). (D) immunostaining on sections shows complete fusion at the WT OFM midline (thick arrow) with laminin-rich BM at the inner limiting membrane (open arrows). In contrast NTN1^-/-^ pioneer cells have not mixed across the midline and retain enrichment for F-actin (arrowheads) and maintenance of their folding points (DAPI, asterisks). However, the BM had remodelled in NTN1^-/-^ (arrows). (E) At HH40 WT OFMs show typical retinal lamination, but NTN1^-/-^ fissure margins were folded with overgrowth of pseudo-laminated structures. The junction of RPE and pioneer cell region is retained in these fissures (small arrows and the folded overgrowth appears attached to the RPE. (F) Overgrown NTN1^-/-^ fissure tissue was positive for outer (ONL) and inner nuclear layer (INL) neural retina markers VAX1 and OTX2. (G-H) Folded overgrowths were also positive for ganglion cell (GCL) and inner nuclear layer (INL) nuclear markers PAX6 and SOX2 (arrowheads) and indicated a broadly laminated structure. S.B. = 50 µm.

### Netrin-1 loss is associated with enrichment for pioneer cell cadherins

To further investigate the lack of cell mixing in netrin-1 mutant OFMs, we stained for N-cadherin and E-cadherin. These adhesion molecules have been previously shown to display reciprocal localisation in pioneer cells and define the boundaries of neural retina and RPE (Hardy and Rainger, 2023). In WT fissures, E-cadherin is specific for RPE and extends at low levels into pioneer cells, whereas N-cadherin is neural retina specific and only extends into pioneer cell regions at low levels (Fig.3A and D). In contrast, we found a trend for cadherin enrichment in mutant fissures. N-cadherin levels were low at the edges of WT pioneer cells (Fig.3A), whereas it was enriched in the same location in mutant fissures. Quantification of N-cadherin signal detected across the full proximal-distal axes showed there was general N-cadherin enrichment in mutants compared to WT (Fig.3B) and that this was significant within 20 µm before and after the fusion point (Fig.3C). Enrichment was also seen in mutants compared to WT for E-cadherin (Fig.3D-F), where detectable E-cad extended more strongly into the pioneer cells at the edges of the margins for the entire OFM. This data shows that loss of Netrin-1 results in enrichment for cadherin in pioneer cells at the margins, and this may be either the result or the cause of increased cell-cell adhesion and fusion failure.

**Fig. 3.**
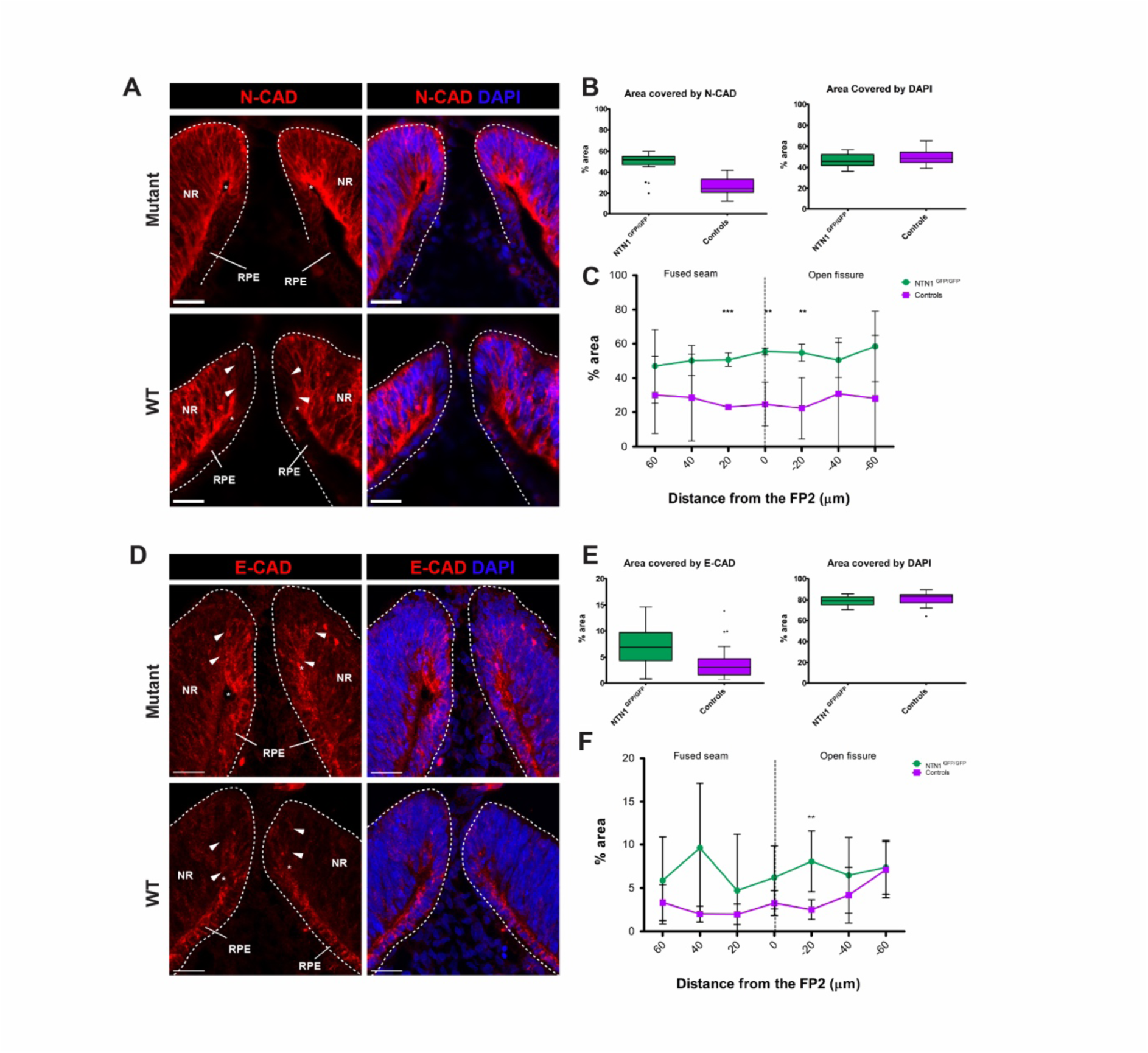
NTN1^-/-^ colobomas associated with increased pioneer cell cadherin. (A) N-cadherin in HH30 OFMs was enriched at the folding point (asterisks) and at the edges of pioneer cells. Intensity plots for total fissure N-CAD (B) and quantification data at key points (C) showed increases in N-cadherin enrichment along the proximal-distal axes of fissures in mutant (NTN1^GFP/GFP^; green) compared to WT (purple). Similarly, immunostaining for E-cadherin (D), intensity plots (E) and quantification data along the proximal-distal axes (F) revealed enrichment in E-cadherin enrichment between mutant and WT embryos. * P<0.05; ** P<0.001; *** P<0.0001; unpaired t-test and f-test to compare variances.

### Reduced apoptosis is a feature of NTN1 mutant optic fissures

To further investigate the consequence of Netrin-1 loss in the OFM, we surveyed ventral eye tissue for apoptosis using antibody staining for activated Caspase-3 (A-Casp-3) in both whole-mount and tissue sections of mutant and WT retinas. We first identified all A-Casp-3 foci in whole-mounted OFMs captured in their full Z-plane (Fig.4A). We collected data for three regions along the fissure margin (including open, fusing and fused fissures), and regions in the peripheral ventral retina not undergoing fusion. The number of foci identified in WT fissures was comparable with previous studies (Hardy et al., 2019) and did not show any difference between WT and mutant fissures in the fused margins, however there was a significant A-Casp-3 reduction in mutant fissures at the fusion point and adjacent open region (Fig.4B). Using sections to distinguish those foci within the retina from the adjacent periocular mesenchyme, we similarly found reduced apoptosis in mutant fissures (NTN1^-/-^) compared to WT (Fig.4C), and this difference was greatest at and around the fusion plate (Fig.4D). In all cases, we did not observe apoptosis in pioneer cells at the immediate margin edges or adjacent to the folding points - foci were generally observed many cell distances away from those cells that directly mediate fusion. These observations were all consistent with our previous findings (Hardy et al., 2019). Thus, loss of Netrin-1 is associated with reduced apoptosis during OFC but is unlikely to be the cause of failed pioneer cell detachment.

**Fig. 4.**
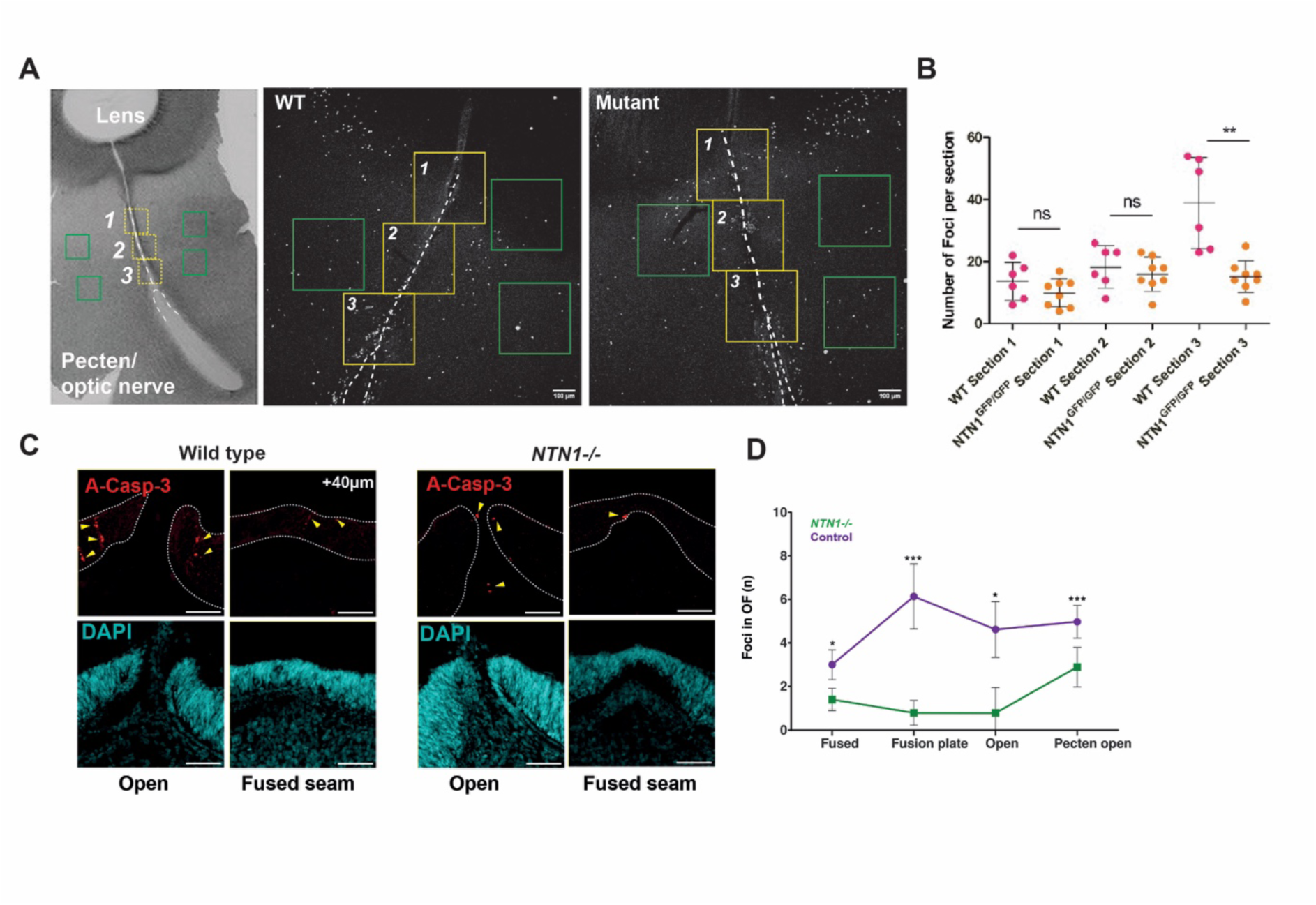
Loss of Netrin-1 correlated with reduced OFM apoptosis. (A) Apoptosing cells identified by whole mount immunostaining of OFM (regions 1-3) and non-fissure regions (green boxes) for A-Casp-3. Hatched line indicates actual OFM. (B) Quantification of A-Casp-3. (C) Representative images of A-Casp-3 on sections for WT and NTN1^-/-^ indicating apoptotic foci (arrowheads). (D) Quantification of apoptotic foci within hatched regions along indicated points in the proximal-distal axes. ns = not significant; * P<0.05, ** P<0.001 and *** P<0.0001. ns = not significant.

### BM remodelling exposed pioneer cells to periocular mesenchyme

Our observation that the BM was correctly remodelled in the NTN1^-/-^ OFMs prompted us to examine the dynamics of BM in normal OFC. Co-staining for BM markers Laminin and Netrin-1 during active fusion revealed that BM dissolution preceded epithelial reorganisation and appeared to transiently expose pioneer cells at the margin edges to the adjacent periocular mesenchyme (Fig.5A; Suppl.Fig.S3). We analysed the previous literature to see if this had been noted previously and found that in mouse, BM removal from the OFM margins also appeared to expose cells to periocular mesenchyme or endothelial cells (Hero, 1989), and similar observations had since been made in zebrafish OFC (James et al., 2016). The significance of these previous findings had not been determined, however. In addition, mutations in several genes expressed in the POM cells have been previously associated with coloboma (reviewed in Patel and Sowden, 2019). Thus, we explored this observation further to ask whether inductive cues may exist in the POM that could influence pioneer cell behaviours.

**Fig. 5.**
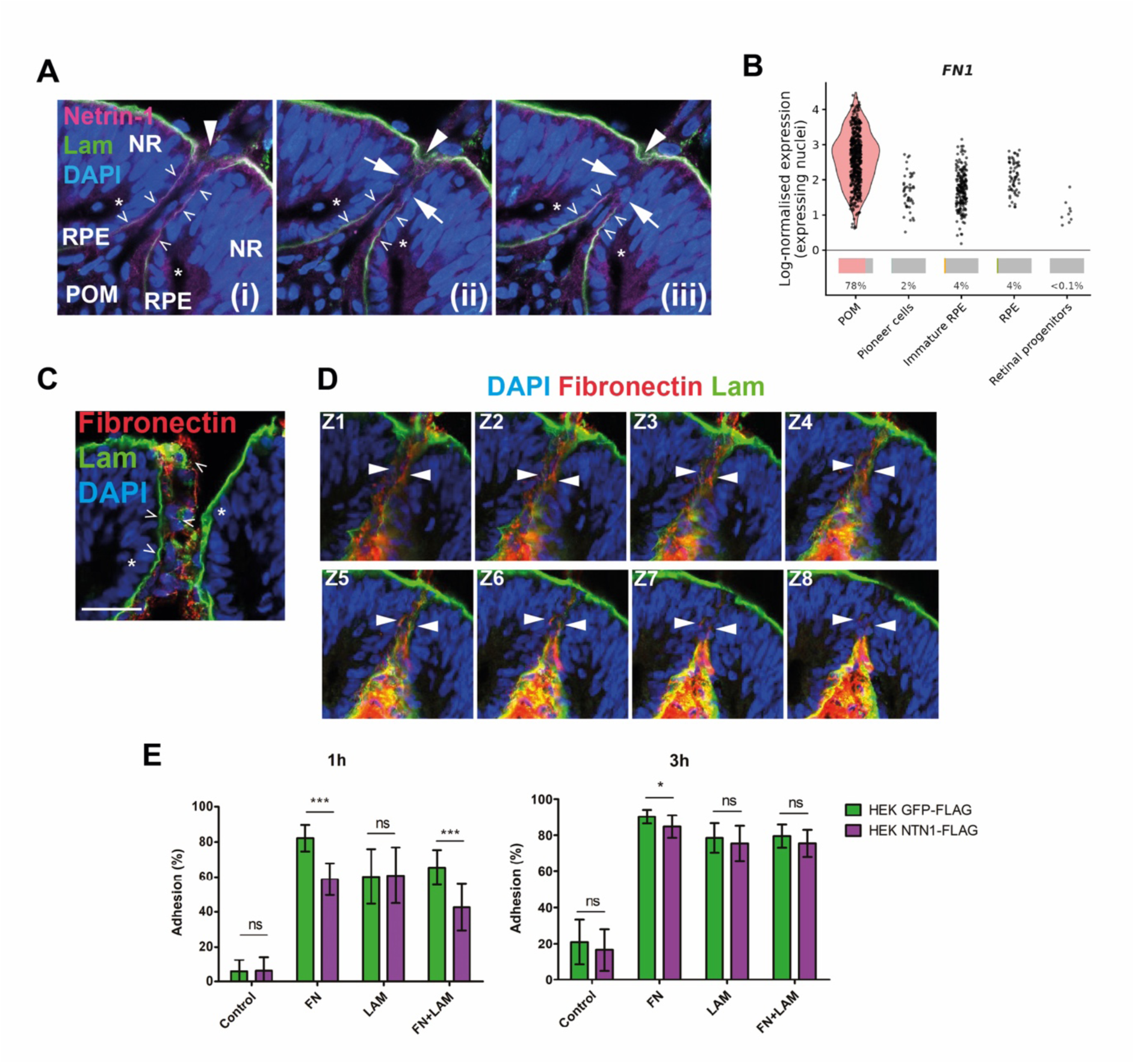
Evidence for BM breakdown exposure of pioneer cells to POM derived Fibronectin. (A) Confocal Z-slices for Netrin-1 and Laminin. (i) Laminin and Netrin-1 containing BM (open arrowheads) initially separate pioneer cells from POM. (ii) Partial BM removal (arrows). (iii) Complete BM removal (arrows). Note the remodelling of the interior BM (large arrowheads), and maintenance of the folding point (asterisks). (B) snRNAseq of WT HH30 OFMs - five cell group clusters are shown. FN1 was detected in 78% of POM cluster, but under 5% of nuclei in every other cluster. (C) POM derived Fibronectin is separated from pioneer by the laminin containing BM. (D) Confocal Z-slices (sequential Z-slices of 1 µm increments) indicated that fibronectin mixed with laminin fragments during BM breakdown. Both are directly localised at pioneer cell domains prior to cell mixing (as indicated by folding point, arrowheads). S.B. = 50 µm. (E) Adhesion assays using Netrin-1 or GFP expressing Flp-In HEK293 inducible cells showed that laminin (LAM) and fibronectin (FN) both increased cell adhesion after 1 and 3h, compared to BSA coated controls. Netrin-1 decreased adhesion on fibronectin-coated plates, and with fibronectin and laminin compared to GFP after 1h. No adhesion effect for Netrin-1 was observed with laminin-only coated plates, and differences were largely resolved by 3h after plating (* P<0.05; *** P< 0.0001).

### Evidence for pioneer cell interactions with POM derived factors

We performed single-nuclei RNAseq (snRNAseq) profiling of the WT chicken OFMs to uncover gene candidates and plausible molecular interactions between the POM and pioneer cells. We first identified the pioneer cell and other cell-type clusters in the snRNAseq data, as defined by previous OFM profiling (Chan et al., 2025; Trejo-Reveles et al., 2023, Patel et al., 2020) (Suppl.Fig.S2). Our analysis showed that FN1 (encoding fibronectin) was selectively and highly expressed in POM nuclei (78% of POM nuclei; Fig.5B) and was rarely detected in pioneer cells (<5% in all other clusters), RPE, or retinal progenitors. Fibronectin is a potent activator of EMT in epithelial cells and has been shown to directly and indirectly interact with Netrin-1 (Brown et al., 2011; Lee et al., 2008; Park and Schwarzbauer, 2014; Yebra et al., 2003). Using immunofluorescence on WT fissures, fibronectin protein was detectable in the POM prior to BM modulation and was clearly separated from pioneer cells by the BM before active fusion begins (Fig.5C). However, during active fusion and BM remodelling we were able to detect regions where fibronectin was interspersed with laminin fragments and appeared to be co-localised with pioneer cells at the fissure margin edges (Fig.5C). To assess whether Fibronectin and Netrin-1 may interact molecularly, we then performed cell adhesion assays using Flp-In HEK293 cells and different ECM substrates (Fig.5D). Attachment of Netrin-1 or GFP expressing cells was no different in plates coated with BSA at either 1hr or 3h, however plates pre-treated with fibronectin, laminin, or a combination of laminin and fibronectin all showed increased cell adhesion in both Netrin-1 and control (GFP) cells at both time points. However, this adhesion was significantly reduced in the combined presence of Netrin-1 with fibronectin at 1h and was more pronounced for Netrin-1 in combination with fibronectin and laminin at 1h. No difference between Netrin-1 or GFP expressing cells was detected on laminin alone. These data show that Netrin-1 can interact with fibronectin to inhibit cell adhesion, and this effect is more pronounced when fibronectin is combined with laminin but does not appear to occur with laminin alone. These data suggest that in WT OFMs, pioneer cells are transiently exposed to both fibronectin and laminin fragments immediately prior to epithelial remodelling, and these interactions (possibly with other POM derived factors) could mediate the cell adhesion changes required to trigger epithelial detachment, and that in the absence of Netrin-1, this mechanism does not occur.

### ITGB8 is a Netrin-1 binding candidate in OFC

As Netrin-1 and Fibronectin are both secreted factors, to evoke pioneer cell responses these would predictably require interactions with receptors at the cell surface. We therefore sought to identify candidate netrin binding partners in pioneer cells using our WT snRNAseq data. Netrin-1 can interact with a range of cell receptors through binding of its specific protein domains (Fig.6A). Of the canonical Netrin-1 receptors, DCC is not detected in the chicken genome (Friocourt et al., 2017), however UNC5 receptors and DSCAM were detected in low numbers of pioneer nuclei, while NEO1 is detected in about half of them but was also broadly present in most retinal cell types and other periocular cell types (Suppl.Fig.S2). It is possible that Netrin-1 could function through NEO1 or other membrane receptors to elicit an adhesive effect, but this would not fit a pioneer cell function requiring spatial and temporally regulated cell detachment at the pioneer cell edges. Integrins are single pass transmembrane proteins that function as alpha and beta heterodimers to physically link the intracellular cytoskeleton to the ECM and adjacent cells, regulating cell adhesion and facilitating changes such as EMT and invasive growth (Harburger and Calderwood, 2009). Integrins such as α6β4 and α3β1 can associate with netrin-1 through their C-terminal domain in several epithelial tissues (Yebra et al., 2003) (Fig.6A) but have not yet been associated with Netrin-1 in fusion (Chaturvedi and Murray, 2021). Integrins have also been shown to mechanistically “zipper” the fusing neural tube via interactions with extracellular fibronectin (Molè et al., 2020), and integrin signalling stimulated by fibronectin can mediate epithelial cell migration and EMT like responses (Brown et al., 2011; Mitra et al., 2011). Thus, we considered integrins as potential OFM Netrin-1 receptors and specifically looked for heterodimers where both pairs were expressed in pioneer cell nuclei. Co-expression for both components of the αV heterodimer pairs ITGAVB8 and ITGAVB5, were detected in pioneer cells; ITGAV was detected in 43% of pioneer nuclei, ITGB8 in 34%, and ITGB5 in 17%. ITGAV was also expressed in the retinal progenitors and POM, but not RPE; whereas ITGB8 was expressed only in pioneer cells and RPE (Fig.6B). Colorimetric In situ hybridisation at prefusion stages (HH28) confirmed that ITGAV was broadly expressed, consistent with our snRNAseq (POM, NR and pioneer cells), whereas ITGB8 was detected in pioneer cells (Fig.6C). Combined fluorescence in situ hybridisation for NTN1 and ITGB8 confirmed their co-expression in pioneer cells prior to fusion (Suppl.Fig.S3) and directly in the opposed OFM margins during active fusion (Fig.6D). Importantly, ITGB8 expression was observed in cells at the very edges of the fissure where epithelial remodelling occurs but was then rapidly reduced in the nascently fused seam, an expression dynamic suggesting a functional requirement for fusion. To then test if Netrin-1 and ITGB8 functionally interact, we performed co-immunoprecipitation on Flp-In HEK293 cells carrying tetracycline inducible Netrin-1:FLAG or GFP:FLAG (Suppl.Fig.S3). We transfected these with a construct expressing a synthesised human ITGβ8 tagged at the C-terminus V5 epitope, and successfully immunoprecipitated Netrin-1:FLAG with ITGB8-V5 (Fig.6E). We did not detect immunoreactivity for control immunoprecipitations between ITGβ8-V5 and GFP:FLAG, or from cells expressing either NTN1:FLAG or ITGβ8-V5 alone. Using A549 cells transfected with ITGβ8-V5 we were also able to co-immunoprecipitate both co-transfected Netrin-1 and Netrin-1 RGDΔ (deletion of cell attachment site) with ITGβ8-V5, but not the GFP control (Fig.6F). In combination, these interaction and expression studies suggest that Netrin-1 can interact with αVβ8 at the protein level and is therefore a plausible receptor in pioneer cells.

**Fig. 6.**
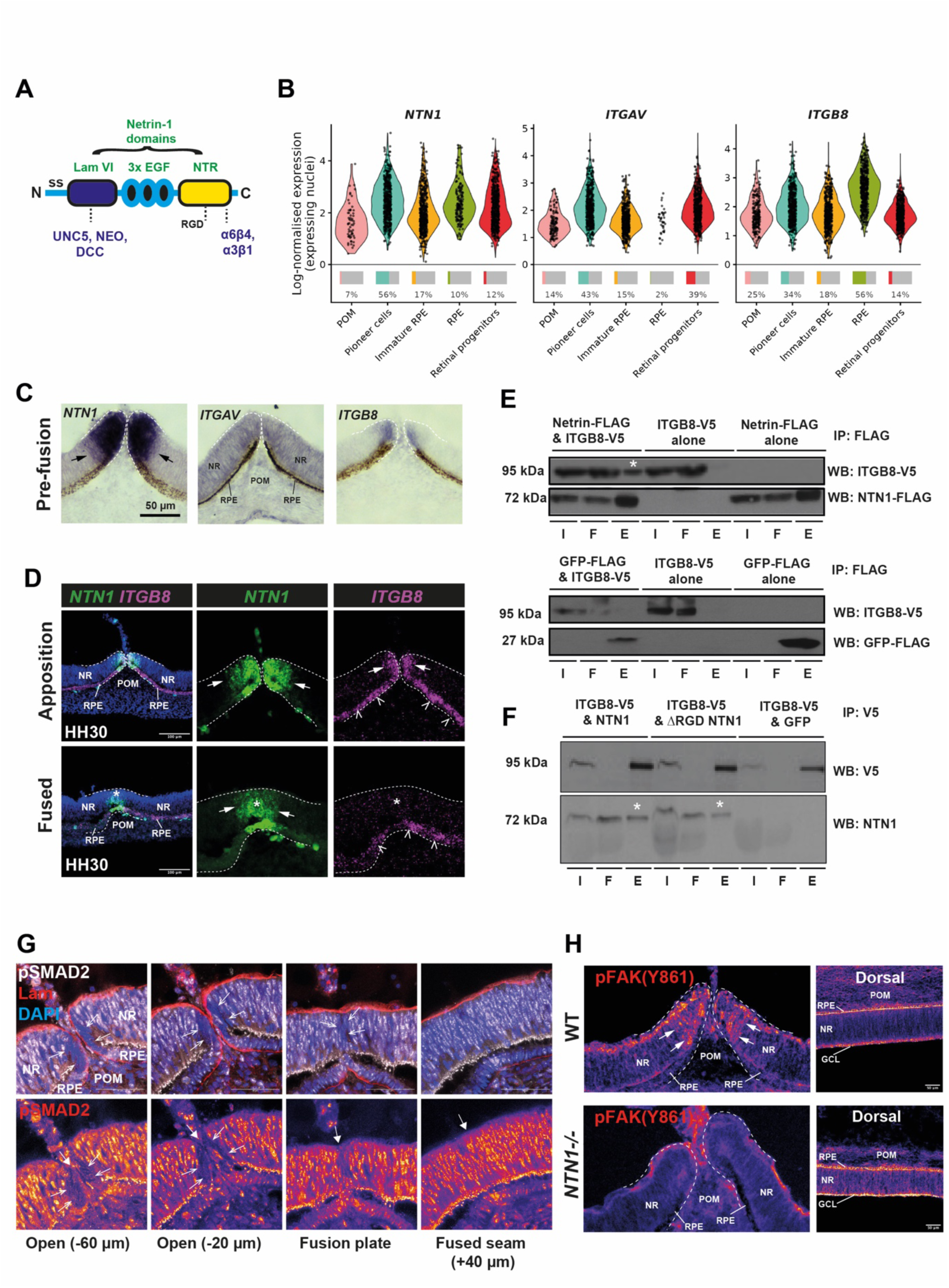
Evidence for Netrin-1 mediated fusion initiation via Integrin signalling. (A) Netrin-1 domains and binding partner interaction sites. (B) Boil plots of snRNAseq data for NTN1, ITGAV and ITGB8. Each gene has its own scale. (C) ITGαV was expressed throughout the POM, NR, and pioneer cell region at HH28 (24h prior to fusion). ITGβ8 was expressed at the fissure margin edges and NTN1 had a broader expression domain. S.B. = 50 µm. (D) Fluorescence in situ hybridisation confirmed co-expression of NTN1 and ITGβ8 in pioneer cells (arrows), and ITGβ8 expression throughout the RPE. (E) Top: Immunoblot of immunoprecipitation with FLAG for NTN1-FLAG cells transfected with ITGB8-V5. Netrin-1 immuno-precipitated with ITGB8 (asterisk). Bottom: immunoprecipitation with FLAG for GFP-FLAG expressing cells transfected with ITGB8-V5 did not immuno-precipitate ITGB8. Key: I, lysates, F, flow-through; E, eluted. (F) Co-immunoprecipitations for A549 cells transfected with ITGB8-V5 and WT or RGDΔ Netrin-1 showed positive interactions between both Netrin-1 versions and ITGB8-V5, but not for ITGB8-V5 in GFP controls. (G) Laminin and phosphorylated SMAD2 immunofluorescence in WT HH30 OFMS during active fusion (n=3). Pioneer cells were absent for pSMAD2 positive nuclear staining at the margin edges, whereas positive nuclei were observed throughout the ventral neural retina, POM and some RPE cells. Whole arrows indicate the OFM midline. Fire LUT panels are shown for pSMAD2 signal only (lower row). S.B. = 50 µm. (H) Phosphorylated FAK (Tyr861) showed enrichment in WT but not NTN1^-/-^ pioneer cells, whereas ganglion cell layers and RPE in dorsal retinas were equivalent between genotypes. S.B. = 50 µm. Fire LUT used to help visualise differences.

### Downstream integrin signalling activity in pioneer cells

Integrins can initiate intracellular responses via phosphorylation and nuclear translocation of SMAD2/3, or through phosphorylation and activation of focal adhesion kinases (FAK) (Deng et al., 2024; Harburger and Calderwood, 2009). To determine if integrin signalling was active in the fissure margin, we performed immunofluorescence for phosphorylated SMAD2 (pSMAD2) in WT fissures (Fig.6G). We found that pSMAD2 was present in nuclei throughout the ventral eye, in the neural retina, in some RPE, and in POM cells. However, nuclear pSMAD2 was absent from fissure margin pioneer cells in the open fissure and in the active fusion plate, indicating that active TGFß signalling is not a direct feature of pioneer cell function in normal OFC. In the fused seam (60 µm after fusion), nuclear pSMAD2 was detected in nuclei at the midline, indicating there is active TGFß signalling in these cells after fusion is complete. Therefore, TGFß signalling is not likely to be a direct outcome of active αVβ8 signalling in pioneer cells unlikely to activate epithelial remodelling during OFC. We then assessed phosphorylated focal adhesion kinase at Tyrosine 861, an indicator of active integrin signalling for EMT responses in epithelial cells (Nakamura et al., 2001). We observed enrichment in RPE and GLC cells of dorsal retinas in both WT and NTN1^-/-^ at HH30. Enrichment of pFAK(861) was also detected in the pioneer cell region in WT (Fig.6H), but we could not detect pFAK(861) reactivity in pioneer cells in NTN1^-/-^ samples at the same stage (n=3 eyes tested per genotype). This data suggests integrin may activate intracellular pFAK signalling in pioneer cells for fusion through Netrin-1. Additionally, signals from the POM that are transiently available to pioneer cells during BM modulation may influence this interaction further to stimulate cell dissociation required for active fusion.

## DISCUSSION

Our Netrin-1 colobomas differ from previously reported phenotypes. Failed OFC can result from persistence of the BM over the fissure margins, as in Fgfr1/2 mutant (Cai et al., 2013; Chen et al., 2012) and Vitamin A deficient mice (See and Clagett-Dame, 2009), or from insufficient ventral optic cup growth that prevents the margins meeting, as in SALL2-associated colobomas (Kelberman et al., 2014). Neither mechanism was observed in our Netrin-1 mutants: eye size was normal, the margins met, and the BM was remodelled normally, so our data show that Netrin-1 is not required for growth of the optic cup or BM remodelling. This apparently differs from NTN1^-/-^ mice where eye size is normal, but BMs appeared to not be remodelled (Hardy et al., 2019). This may be species specific, or reflect the developmental stage analysed in those mice (E11.5), when OFC is partially complete. In our mutant chickens, pioneer cells maintained their apical F-actin at the folding points of the fissure and did not mix across the midline, despite removal of the overlying BM, leaving a folded overgrowth of ventral neural retina and persistent coloboma. This indicates that BM remodelling and pioneer cell detachment can be mechanistically separated, and that detachment depends on Netrin-1. The closest reported phenotype is in Shroom3 mutant mice, which also develop folded retinal epithelium (Herstine et al., 2025). There, the BM was remodelled and cells detached and mixed, but apical-basal polarity was not re-established, leaving a thinner, folded neural retina at the fusion point.

### Netrin-1 interactions in OFC

Our data suggest that Netrin-1 could functionally engage integrin αvβ8 during OFC, however we could not confirm this interaction in vivo or show a functional requirement for αvβ8. Netrin-1 co-purified with ITGB8 and the interaction was retained with an RGD-mutant Netrin-1, consistent with binding through the C-terminal region rather than the RGD cell-attachment motif (Yebra et al., 2003). ITGB8 is considered non-canonical because its cytoplasmic tail diverges from that of other β-integrins, and most evidence supports a role in activating latent TGFβ at the cell surface (Mu et al., 2002). However, pSMAD2 was selectively absent from pioneer cell nuclei in our analyses, arguing against canonical TGFβ activation as the ITGB8 effector. ITGB8 knockdown reduces FAK phosphorylation in endometrial epithelial cells (Kumar et al., 2017), so we asked whether ITGB8 instead engages focal adhesion signalling. pFAK(Y861) was enriched in pioneer cells but absent in NTN1^-/-^ eyes. Existing RNAseq data show ITGB8 is expressed in the optic fissure of humans, mice and zebrafish (Chan et al., 2025; Hardy et al., 2019; Richardson et al., 2019). Itgb8 mutants have not been screened for coloboma, but cleft palate in Itgb8 and Itgav mutants (Aluwihare et al., 2009; Bader et al., 1998) hints at roles in other fusion contexts, although in the palate this has been attributed to loss of TGFβ3 activation. Direct analyses of ITGB8 function in OFC and ocular phenotyping in existing genetic models are needed to confirm its role in pioneer cell detachment.

### Netrin-1 in pioneer cell detachment

In NTN1^-/-^ embryos, removal of the BM brought pioneer cells from the two margins into direct apposition but did not trigger epithelial reorganisation, suggesting that detachment needs a separate initiating cue that depends on Netrin-1. NTN1 and ITGB8 are co-expressed at least 24h before fusion (ITGB8 at HH28 here; and NTN1 at ∼HH22 in Hardy et al., 2019), and NTN1 is downregulated immediately after (Hardy et al., 2019; Richardson et al., 2019). We therefore suggest that Netrin-1 primes pioneer cells for detachment rather than executing it, with the trigger coming from the POM or other periocular cells transiently exposed to pioneer cells. We found evidence for an altered adhesive state in mutant pioneer cells: N-cadherin and E-cadherin both showed a localised increase in pioneer cells at the very edge of the fissure compared to WT. Elevated cadherins could cause the detachment failure or result from cells being unable to leave the folding point building up excess cadherin, and our data cannot distinguish these. In other tissues Netrin-1 promotes adhesion, for example in the mammary gland via neogenin (Srinivasan et al., 2003) and in pancreatic epithelium via α6β4 and α3β1 integrins (Yebra et al., 2003). Our finding that Netrin-1 reduced adhesion to fibronectin points towards a context-dependent effect, reflecting cell type, matrix composition or receptor interactions.

### Towards a mechanistic role for POM in OFC

Mutations in POM-expressed genes and experimental removal of POM cause colobomas (Patel and Sowden, 2019; Gestri et al., 2018), but the specific requirement for the POM in OFC is unknown. Periocular phagocytic cells in the mouse fissure (Hero, 1989) and endothelial cells in zebrafish fissures (James et al., 2016) were seen adjacent to exposed pioneer cells at BM breakdown, but these were discussed in the context of debris clearance or BM dissolution, not communication with pioneer cells. We found fibronectin, whose expression appeared restricted to the POM, together with laminin from the overlying BM, co-localised as fragments at the exposed pioneer cell regions as the BM broke down. This is consistent with pioneer-periocular cell contact at the point of fusion, and marks fibronectin as a strong candidate cell-detachment cue. A direct Netrin-1-fibronectin interaction was previously confirmed (Yebra et al., 2003), and Netrin-1 can act as a substrate-bound (haptotactic) cue that modifies cell responses to fibronectin (Lee et al., 2008; Yebra et al., 2003), so a functional interaction at the pioneer-POM interface is highly plausible. Fibronectin may act as a ligand triggering intra-cellular signalling in pioneer cells, or as a mechanical substrate exerting traction on them to physically move across the midline. The reduced fibronectin adhesion we see with Netrin-1 in our cell assays is easier to reconcile with a signalling model than with pure traction. Selective removal of fibronectin in periocular cells at the fissure, or targeted interference to fibronectin-receptor interactions would show whether it is required for OFC, and by what mechanism.

### Relevance to human coloboma

A heterozygous de novo missense variant, NTN1 p.(Tyr495Asn), was recently reported in a patient with unilateral chorioretinal coloboma, microphthalmia, sensorineural hearing loss and polydactyly (Toms et al., 2025). Because heterozygous loss of Netrin-1 does not cause colobomas in chicken or mouse (Chan et al., 2025; Hardy et al., 2019), a dominant-negative or gain-of-function effect may be more plausible than simple haploinsufficiency, although humans may be more dose sensitive: heterozygous NTN1 variants that reduce Netrin-1 secretion cause congenital mirror movements (CMM; Méneret et al., 2017), a condition also seen with DCC mutations (Srour et al., 2010). The functional effect of the coloboma NTN1 variant is unknown, but would be consistent with altered integrin binding, since integrins bind the C-terminal region of Netrin-1 (Yebra et al., 2003), whereas Netrin-1 interactions with DCC are mediated by the laminin and EGF domain (Finci et al., 2014; Xu et al., 2014). With no other variants so far identified in coloboma patients, causality cannot yet be confirmed. Netrin-1 may also act in the human inner ear: the patient had hearing loss, and ntn1a zebrafish morphants lose sensory hair cells (Toms et al., 2025). We did not examine the chicken cochlea, but the semicircular canals, which fail to fuse in NTN1 mutant mice (Salminen et al., 2000), appeared grossly normal in our knockouts. This may reflect compensation by NTN2 during fusion, which is co-expressed with NTN1 in the chicken canal fusion plate (Nishitani et al., 2017), but is not detected in vertebrate OFMs (Chan et al., 2025; Hardy et al., 2019; Richardson et al., 2019) and so may leave the eye more sensitive to NTN1 loss compared to the fusing inner ear.

### Apoptosis in OFC

Apoptosis, which occurs during normal chick OFC (Hardy et al., 2019), was reduced in NTN1 mutant OFMs, most strongly at the fusion point and adjacent open region but not in the fused margin. The requirement for cell death in OFC is controversial and species-specific: chicken and mouse OFMs contain apoptotic cells, whereas zebrafish and human fissures do not (Patel et al., 2020; Chan et al., 2021). In WT chicken OFMs, cell death peaks in the fused seam where Netrin-1 levels have been downregulated, and those foci observed immediately before fusion lie in the wider ventral retina at the boundary of the Netrin-1 domain, rather than directly in pioneer cells (Hardy et al., 2019). These suggest that specific dosage of Netrin-1 may trigger apoptosis in the OFM. Netrin-1 is proposed to promote survival by blocking apoptosis induced by the dependence receptors DCC and UNC5H (Llambi et al., 2001; Mehlen et al., 1998), although this is disputed in vivo (Bin et al., 2015; Yung et al., 2015). DCC is absent from the chicken genome (Friocourt et al., 2017), and we did not detect UNC5 paralogues in pioneer cells. However, loss of a survival factor would also be expected to increase apoptosis whereas we observed a reduction, so reduced apoptosis in NTN1^-/-^ fissures is unlikely to reflect loss of dependence receptor signalling. Instead, it may reflect NTN1 expression dynamics or be secondary to failed fusion, and our data cannot distinguish these. The relationship between Netrin-1, apoptosis and fusion also differs in the chicken inner ear, where apoptosis is required for fusion (Fekete et al., 1997) and Netrin-1 overexpression prevents it (Nishitani et al., 2017). Whether apoptosis contributes to pioneer cell removal in the chicken eye, or follows aberrant cell behaviours during and after fusion, remains untested.

### Germline chicken models for developmental biology and disease

The Netrin-1 knockout chicken, to our knowledge the first gene-edited avian model with a developmental eye phenotype, highlights the value of this approach for developmental biology applications (McGrew et al., 2025). It demonstrates that germline editing in the chicken can model a developmental disorder and suggests this could be extended to test the causality of human DNA variants by screening allele-specific, gene-edited PGC-derived embryos for phenotypes and investigating their underlying molecular mechanisms. Indeed, specific point mutations have been previously generated in chickens (Idoko-Akoh et al., 2023), and this could be extended to generate human disease specific variants. Furthermore, pooling independent PGC clones carrying different variants through one surrogate host could theoretically allow even multiple variants to be tested simultaneously in progeny, while editing multiple loci in the same clones could test digenic or oligogenic contributions. Our work therefore provides a timely foundation for using germline chicken models to link disease-associated genes to developmental disorders and their mechanisms.

## METHODS

### Generation of NTN1 knockout PGCs

We used the same NTN1 gene editing strategy in PGCs as used previously to generate the NTN1-T2A-eGFP line (Chan et al., 2025), with the exception that we did not co-transfect a repair template into PGCs. Briefly, male PGCs were isolated from day 2.5 Hy-Line chick embryonic blood samples and selectively grown using previously published protocols (Whyte et al., 2015). PGCs were transfected using Lipofectamine 2000 (Thermo Fisher Scientific) according to the manufacturer’s instructions with the CRISPR/Cas9 plasmid pX458 (Ran et al., 2013) containing the NTN1-specific sgRNA sequence: 5’-GACCCCTGCTACGACGAGCA-3’. Transfected PGCs were FACS sorted to seed individual cells into 96-well plates following the protocol of Ballantyne et al. (Ballantyne et al., 2021). Of 120 single cell cultures, we eliminated dead clones as well as the fastest and slowest growing clones. For the remaining clones, we cryopreserved cell stocks for injections and screened the clonal populations by genomic DNA extraction and sequence specific PCR, using oligonucleotide primers specific for the endogenous NTN1 (exon 1) locus (Table 1).

**Table 1.**
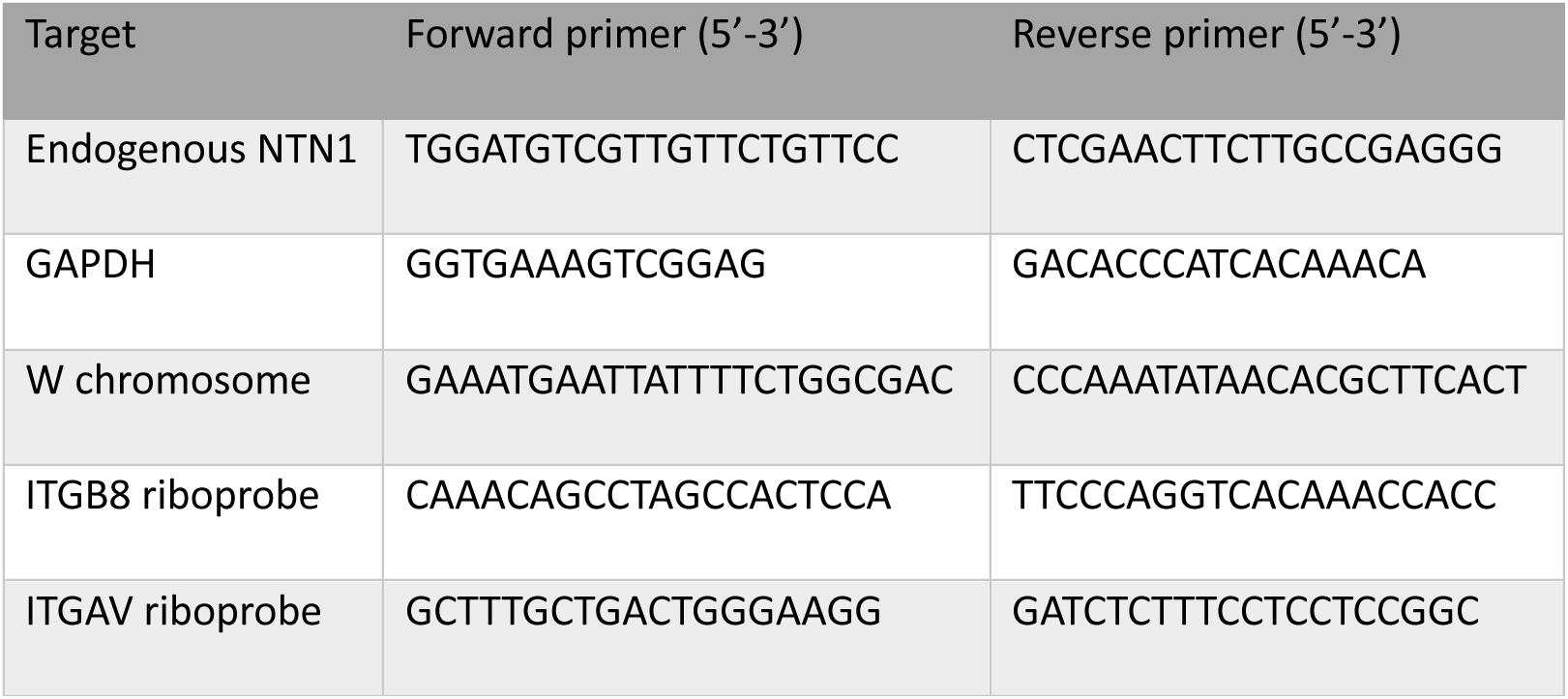
Oligonucleotide primers used for PCR based genotyping and qRT-PCR.

### Generation of a NTN1 knockout chicken line

Chicken lines were generated at the National Avian Research Facility, Roslin Institute. All animal management, maintenance and embryo manipulations were carried out under UK Home Office licences PP0018931 and PP1196147. Experimental protocols and studies were approved by the Roslin Institute Animal Welfare and Ethical Review Board Committee (AWERB references #P271 and #P294) and were performed in accordance with the relevant approved guidelines and regulations therein. All data and analyses in the study are reported in accordance with ARRIVE guidelines (https://arriveguidelines.org). A clonal male targeted PGC line was thawed and injected into iCaspase9 surrogate host embryos according to previously published protocols (Ballantyne et al., 2021). Briefly, to remove endogenous germ cells in the host chicken embryos 1.0 µl of 25 mM B/B (in DMSO) (Takara Bio) was added to 50 µL PGC suspensions before injection. Chicks (n=4/12) were hatched, and males (4/4 G0 birds hatched) were raised to sexual maturity. G0 males were placed in breeding pens with 6-7 WT Hy-Line female layers and eggs collected on a weekly basis. Eggs were incubated at 37.5°C up to 14 days or hatched for breeding colonies. G1, G2 and G3 generations of heterozygous birds were co-housed to collect offspring of all genotypes. No offspring from heterozygous matings were taken past embryonic day 14 or hatched in this study.

### Genotyping

Tissues were collected from embryos, and genomic DNA was extracted using the PureLink Genomic DNA Mini Kit (Thermo Fisher Scientific) according to the manufacturer’s instructions. PCR and Sanger sequencing was performed using NTN1 specific primers (Table 1) to confirm inheritance of the NTN1 knockout alleles, and sex typing used W chromosome specific primers. To test whether the NTN1 knockout allele was transmitted at the expected Mendelian ratio, embryos from heterozygote × heterozygote crosses were genotyped (n=937). The observed numbers of WT, heterozygous and homozygous embryos were compared with the expected 1:2:1 ratio using a Pearson χ² goodness-of-fit test with 2 degrees of freedom. Significance was set at p< 0.05. Primers for genotyping NTN1^T2A-eGFP^ are as previously used (Chan et al., 2025).

### Preparation of samples for western blotting

Whole optic fissures or neural tube tissues were microdissected dissected from HH28 NTN1^T2A-eGFP/+^ embryos into ice cold lysis buffer (50 mM Tris (pH 7.4), 150 mM NaCl, 1% NP-40, 0.5% sodium deoxycolate, 0.1% sodium dodecyl sulfate, supplemented with Roche cOmplete EDTA-free protease inhibitor cocktail immediately prior to use), mechanically homogenised using a microtube pestle and incubated on ice for 30 minutes with occasional vortexing. Samples were then clarified by centrifugation (12,000 x g for 30 minutes at 4 °C) and protein concentration estimated by BCA microplate assay before western blotting. Adherent cell line samples had media removed and two gentle warm PBS washes prior to ice cold lysis buffer being added directly to the cells. Samples were then kept on ice with gentle agitation for 15 minutes before being scraped into a microfuge tube and clarified as described above. Protein concentration was then estimated by BCA microplate assay before western blotting.

### Western Blotting

Equal amounts of protein were mixed with 1x NuPAGE LDS sample buffer and 1x NuPAGE sample reducing agent and resolved by SDS-PAGE using discontinuous polyacrylamide gels (4% stacking gel and 10% resolving gel prepared from 30% acrylamide/bis-acrylamide, 29:1). Electrophoresis was performed in Tris-glycine-SDS running buffer (25 mM Tris, 190 mM glycine, 0.1% SDS, pH 8.3) at a constant voltage of 70 V for 20 min to allow proteins to stack and enter the resolving gel, followed by 150 V for 70 min until the dye front reached the bottom of the gel. Proteins were transferred onto 0.45 µm nitrocellulose membranes in transfer buffer (25 mM Tris, 190 mM glycine, 0.1% SDS, and 20% methanol) on ice at a constant current of 250 mA for 90 min. Membranes were blocked in 5% (w/v) non-fat skimmed milk in TBS-Tween20 (v/v 0.1%) with gentle agitation at room temperature for 1 h, followed by overnight incubation at 4 °C with primary antibodies (Table 2) diluted in 5% (w/v) non-fat skimmed milk in TBS-T. Membranes were washed three times for 10 min each in TBST with gentle agitation and incubated with the appropriate HRP-conjugated secondary antibodies (see table) diluted in 5% (w/v) non-fat skimmed milk in TBS-T for 1 h at room temperature. After three additional 10 min washes in TBS-T, immunoreactive bands were visualised using enhanced chemiluminescence (ECL; 32209, Pierce) and X-ray film. Apparent molecular weights were estimated using a pre-stained protein molecular weight marker (#74124, CST).

**Table 2.** Antibodies used for Western Blotting.

| Antibody | Host | Working Concentration |
|---|---|---|
| Primary Antibodies |  |  |
| Anti-Netrin 1 antibody [EPR5428]<br>ab126729, Abcam | Rabbit monoclonal | 0.5 µg/ml |
| ANTI-FLAG M2 antibody<br>F3165, Sigma | Mouse monoclonal | 2 µg/ml |
| V5-Tag antibody [SV5-Pk1]<br>MCA1360GA, BioRad | Mouse monoclonal | 0.5 µg/ml |
| GAPDH (D16H11) XP<br>5174S, CST | Rabbit monoclonal | 1:2000 |
| Anti-GFP [3H9]<br>3H9, Chromotek | Rat monoclonal | 1 µg/ml |
| Secondary Antibodies |  |  |
| Anti-Mouse IgG, HRP-linked<br>A9044, Sigma | Rabbit polyclonal | 1:5000 |
| Anti-rat IgG, HRP-linked<br>A9037, Sigma | Goat polyclonal | 1:5000 |
| Anti-rabbit, IgG HRP-linked<br>A0545, Sigma | Goat polyclonal | 1:5000 |

### Reverse-transcriptase PCR

Eight eyes (from four embryos) were dissected and pooled for each optic fissure replicate, and two replicates for the neural tube. Total RNA was isolated using Trizol reagent (Ambion, Life Technologies) according to the manufacturer’s instructions and treated with Turbo DNaseI (Thermo Fisher Scientific). RNA quality and integrity were assessed using a nanodrop spectrophotometer (Thermo Fisher Scientific) and standard gel electrophoresis. Total RNA was reverse transcribed using oligo-dT primers with Superscript III (Thermo Fisher Scientific) according to the manufacturer’s instructions. PCR was performed using NTN1 and GAPDH sequence-specific primers (Table 1), and amplicons were run on 1% agarose gels to confirm the amplification. qRT-PCRs were performed using Luna Universal qPCR Master Mix (NEB) and an Agilent Stratagene Mx3000P system with the same primers. Cycle conditions: 95°C ×1 minute, followed by 40 cycles of 95°C ×10 seconds; 60°C ×30 seconds; followed by 1 cycle of 95°C ×1 minute; 55°C ×30 95°C ×30s. We performed 3 technical replicates for each biological replicate. To calculate relative gene expression, we use ratios determined by the Pfaffl Method Ratios (Pfaffl, 2001). We use an unpaired t-test with Welch’s correction to identify expression differences.

### Immunofluorescence

Embryos were collected, staged (Hamburger and Hamilton, 1992) and fixed for 2 hours in 4% Paraformaldehyde in phosphate buffered saline (PBS; pH 7.0), then rinsed in PBS and equilibrated in 10% Sucrose-PBS prior to embedding in Neg-50 (Epredia) and snap-freezing in Iso-pentane at -80°C. For immunofluorescence analysis we used 20 µm cryosections incubated with antibodies against Netrin-1 (#AB126729; Abcam; 1:200 dilution), Laminin (#3H11; DSHB; 1:20), GFP (#ab5450; Abcam; 1:500), Phos-SMAD2 (#zrb04953; Merck; 1:200 with 3 day incubation), Phos (Tyr861) FAK (#44-626G; Thermo Fisher Scientific; 1:250), SOX2 (PCRP-SOX2-1B3; DSHB; 1:5) PAX6 (DSHB; 1:5) Fibronectin (#B3/D6; DSHB; 1:50), Laminin (rabbit polyclonal #L9393; Merck; 1:200;), NF145 (#MAB1621; Millipore; 1:100), ZO-1 (#402200; Thermo Fisher Scientific; 1:500), E-Cadherin (#610181; BD Biosciences; 1:200); N-Cadherin (Merck; #C3865; 1:200), overnight at 4°C in 0.1% BSA-PBS with 0.1% Tween. Alexa-Fluor fluorescently conjugated secondary antibodies (Thermo Fisher Scientific) were used at 1:1000 dilutions. F-actin (Phalloidin-488 nm; Thermo Fisher Scientific; 1:200) and DAPI (4ʹ,6-diamidino-2-phenylindole; Thermo Fisher Scientific; 1:1000) were also used as fluorescent counter stains. Slides were mounted with coverslips using Fluorsave (Merck). Whole mount immunofluorescence against activated Caspase-3 on dissected optic fissures was previously described (Hardy et al., 2019). Imaging was performed on Zeiss LSM-800 confocal with Zen Black software (v2.3 Zeiss) for acquisition, or a Leica DMLB upright Fluorescence microscope with Micromanager (v1.4). Images were processed using FIJI-ImageJ (Schindelin et al., 2012).

### Image quantitation

Apoptosis was quantified by counting active caspase-3 (A-Casp3) positive foci in flat-mounted ventral eyes and in cryosections. In flat-mounts, foci were counted in three 200 × 200-pixel region of interest (ROIs) [area, µm²] placed along the OFM, and in three randomly positioned ROIs in non-optic fissure regions, selected with G. Landini’s ImageJ macro RandomSampleAndMeasure. The optic fissure was divided into distal (region 1), central (region 2) and proximal (region 3) regions. Foci were counted with ImageJ’s automatic particle analysis, using a minimum size of 2 µm² to exclude background and particles too small to resolve; the counted foci therefore include apoptotic bodies (78–1963 µm²) and larger microvesicles (Santavanond et al., 2021; Van der Meeren et al., 2023). To exclude secondary-antibody aggregates that can mimic large apoptotic bodies, extreme values were removed with Grubbs’ test (α = 0.05). In cryosections, all A-Casp3 positive foci were counted manually within a fixed 300 × 300 pixel ROI centred on the OFM folding point. Foci in the neural retina (NR) inside the ROI were classed as optic fissure foci. Foci in the POM inside the ROI; and all foci in the NR, retinal pigment epithelium (RPE) and POM outside it, were classed as non-optic fissure foci. Normality was assessed with the Shapiro–Wilk test, and groups were compared with the Kruskal–Wallis test followed by Dunn’s multiple comparison test. WT n=6 OFMs; NTN1^GFP/GFP^ n=8. Eyes from WT and NTN1 homozygous embryos were collected at HH30 and analysed for N-cadherin (N-Cad/CDH2) and E-cadherin (E-Cad/CDH1) by immunofluorescence, with DAPI as a nuclear counterstain. N-Cad was analysed in WT (n=4) and NTN1 mutant (n=4) embryos, and E-Cad in WT (n=3) and NTN1 mutant (n=3) embryos. Serial sections through the optic fissure were imaged and the percentage of area covered by N-Cad or E-Cad staining was measured at defined positions along the fissure, relative to the location of fusion point ±20 µm. The distribution of the N-Cad, E-Cad and DAPI data was examined with Tukey boxplots (box, 25th–75th percentile; line, median) to identify outliers. WT and mutant embryos were compared at each position with an unpaired t-test, with an F test to compare variances. Significance was set at * P ≤ 0.05, ** P ≤ 0.01 and *** P ≤ 0.001.

### In situ hybridisation

Cryosections were prepared for RNAscope as for immunofluorescence, with the exception that slides were air dried for 2h at room temperature, heated to 50°C for 30 mins, then immersed in 70% EtOH for 10 mins, prior to following the RNAscope Multiplex Fluorescent v2 Kit (ACD, Biotechne) protocol for frozen tissue. RNAscope probes were used for ITGB8 (Gg-ITGB8 #1667111-C1 targeting XM_046933080.1), OTX2 (Gg-OTX2 #902291-C2 targeting NM_204520.2 20ZZ), and VAX1 (Gg-VAX1 #1144181-C1 targeting NM_204799.3). For colorimetric in situ hybridisation, we followed a standard protocol described in J. Rainger’s doctoral thesis (available on request) using riboprobes designed for NTN1 (Hardy et al., 2019), ITGAV and ITGB8 (Table 1).

### Gross phenotypic assessments of NTN1 knockout embryos

Embryos were assessed for grossly visible anomalies by brightfield stereomicroscopy. Colobomas were detected at E9, and a subset were analysed for laterality. For eye size analyses, the horizontal and vertical (coronally) dimensions of the eye globe were measured using digital callipers (resolution; 0.01 mm). Horizontal and vertical measurements were averaged per eye, and the average of both left and right eyes were used in comparison analysis. GraphPad Prism was used to perform One-Way ANOVA on datasets. The overall association between genotype and eye defect was tested by χ² test (χ² = 168.2, df = 2, p = 3.1 × 10⁻³⁷). Pairwise Fisher’s exact tests on total eye defect with Holm correction for three comparisons was performed (HOM vs WT, p = 3.9 × 10⁻²⁴; HOM vs HET, p = 1.3 × 10⁻²⁹; WT vs HET, not significant, ns). **** p< 0.0001. Laterality was scored in an independent cohort of NTN1^-/-^ embryos (n=35 embryos with coloboma) and is reported descriptively as the number and percentage of affected embryos in each class. No statistical tests were applied to these data.

### Soft tissue micro-Computed Tomography

HH34 embryos were prepared for µCT using 0.1M phosphotungstic acid at 40°C for 48h as previously described (Naïja et al., 2024; Rzhepakovsky et al., 2023). We used a NeoScan N80 microCT and scanned samples with X-ray voltage 75 kV, X-ray current 200 µA with filter set at Al 1.0 mm, 4.5 μm pixel size, 0.2 degree rotation step, 4x averaging. Data analysis, reconstructions, and volume rendering was performed using NeoScan_80 software version 3.1.2 (13) and FIJI-ImageJ. Inner ear 3D reconstructions were carried out using methodology described in (Bryant et al., 2021). The datasets were statistically analysed in GraphPad Prism using Two-Way ANOVA with Bonferroni post-hoc test.

### Cell spreading assay

A cell spreading assay using inducible NTN1-FLAG and GFP-FLAG Flp-In HEK293 cells was performed to investigate cell adhesion. Plates were coated with various extracellular matrix proteins. Briefly, 12-and 24-well culture plates were incubated with 1% BSA in PBS, fibronectin (#341635 Millipore; 10 μg/ml), laminin (Sigma #L2020; 10 μg/ml), and a combination of fibronectin and laminin (10 μg/ml) and left overnight at 4°C. Cells were dissociated after a brief exposure to 0.25% trypsin-EDTA, then left to recover in fresh growth media with tetracycline for 1h with manual shaking every 10 minutes. Cells were then plated on the different adhesion substrates (4×10^4^ or 8×10^4^cells/well), and attached cells were photographed after 1h and 3h. Images were taken with an Axiovert 5 microscope (Zeiss), and total adhered cells (more than 200 cells per experiment) were measured from randomly selected ROIs (400 μm^2^) using an in-house macro in ImageJ. Data was analysed in GraphPad Prism using Mann Whitney test and 4 biological replicates with 3 technical replicates per experimental group were performed.

### Single nuclei RNAseq

Optic fissures were dissected from HH30 as previously described and were pooled to contain from independent embryos; each pool contained 10 optic fissures from 10 separate embryos. 4 independent pools were prepared on separate dissection days and were snap frozen at -80°C. Samples were then processed using Chromium Nuclei Isolation with RNase Inhibitor Kit (1000494) according to the manufacturer’s instructions at the University of Newcastle Genomics Core. The four isolated single nuclei preps underwent single-nuclei 3’ mRNAseq library preparation using the 10X Genomics Chromium controller following the Chromium Next GEM Single Cell 3ʹ Reagent Kits v3.1 (Dual Index) user guide (CG00315 Rev D). Each library was then QC checked using an Agilent TapeStation and all 4 samples were equimolar pooled. The 4-sample pooled library was then sequenced on a 100 cycle NovaSeq 6000 S1 flow cell.

### Bioinformatics

Each pool’s library was sequenced on two lanes as 28-bp Read 1 (cell barcode and UMI) and 90-bp Read 2 (cDNA), giving 371 to 539 million read pairs per library (1.91 billion in total). Reads were aligned to the chicken GRCg7b assembly (bGalGal1.mat.broiler.GRCg7b, GCA_016699485.1) with the Ensembl release 116 gene annotation, and unique molecular identifiers (UMIs) were counted per gene with Cell Ranger 10.0.0 (10x Genomics; Zheng et al., 2017), including reads in introns, as is appropriate for nuclear RNA. Ambient RNA was removed from each library’s unfiltered count matrix with CellBender 0.3.2 remove-background (Fleming et al., 2023), with 10,000 expected nuclei, 30,000 droplets included and the default target false-positive rate of 0.01; the droplets CellBender called as containing a nucleus, with their ambient-corrected counts, were used from then on. Both steps were run in nf-core/scRNAseq 4.2.0 (Ewels et al., 2020). Genes were identified by their gene symbol from the Cell Ranger feature table, or by Ensembl gene ID where no symbol was annotated (with the ID appended where two genes share a symbol). Barcodes with fewer than 200 UMIs were discarded, and doublets were identified and removed in each library with scDblFinder 1.24.0 (Germain et al., 2021), with the expected number of doublets set from 10,000 recovered nuclei per library. Low-quality nuclei were then removed per library with outlier thresholds of 3 median absolute deviations (MADs) from the library median (Amezquita et al., 2020; Amezquita et al., 2026, quality control chapter): nuclei more than 3 MADs below the median for log-transformed detected genes or UMI counts, or more than 3 MADs above the median for the percentage of UMIs from the 13 mitochondrial protein-coding genes, were removed. The UMI criterion removed no nuclei, as its cut-offs fell below the 200-UMI floor. The gene and mitochondrial thresholds also had absolute bounds: at least 300 detected genes, and an upper mitochondrial threshold bounded between 0.5% and 10%. This left 46,930 nuclei (10,637 to 12,667 per pool). Downstream analysis used Seurat 5.5.1 (Hao et al., 2024) in R 4.5.3. Counts were normalised with SCTransform v2 (Hafemeister and Satija, 2019; Choudhary and Satija, 2022) using the glmGamPoi backend (Ahlmann-Eltze and Huber, 2021). The percentage of mitochondrial UMIs and the difference between the S-phase and G2/M-phase scores were regressed out; the scores came from Seurat’s CellCycleScoring with its human cell-cycle gene lists (Tirosh et al., 2016) matched to chicken gene symbols. Regressing the difference removes variation between cycle phases while keeping the variation between cycling and non-cycling nuclei. Mitochondrial and cell-cycle genes were also excluded from the variable genes before principal component analysis (30 components). The four pools were integrated with Harmony 2.0.5 (Korsunsky et al., 2019), with pool as the batch variable, and the first 15 Harmony dimensions were used for the shared nearest-neighbour graph, Louvain clustering at resolution 0.4 and the UMAP embedding, giving 13 clusters. Clusters were annotated by comparing their marker genes with published chicken optic fissure and pioneer cell markers (Trejo-Reveles et al., 2023; Chan et al., 2025) and with literature markers of other ocular cell types. Candidate markers were identified with Seurat’s Wilcoxon rank-sum test on SCTransform-corrected expression and ranked by log2 fold change. Each cluster was also compared with the rest of the nuclei by pseudobulk differential expression with DESeq2 1.50.2 (Love et al., 2014): counts were summed per pool inside and outside the cluster and compared with the model ∼ pool + cluster membership, and markers were called at Benjamini-Hochberg adjusted p< 0.05, shrunken log2 fold change ≥ 1 (DESeq2 normal prior) and detection in at least 10% of the cluster’s nuclei. All 46,930 nuclei were used for clustering and marker testing. One cluster (cluster 10; 1,455 nuclei) had a low-quality profile, with low UMI counts and ribosomal protein genes as its top markers and was left out of the gene-detection summaries and the dot plot, which therefore cover 45,475 nuclei in 12 clusters. For plots and per-cluster summaries, counts were log-normalised (Seurat LogNormalize, scale factor 10,000). A gene was counted as detected in a nucleus with at least one UMI. For Figures 5B and 6B, clusters were grouped as periocular mesenchyme (cluster 11), pioneer cells (6), immature RPE (4), RPE (7) and retinal progenitors (0, 1 and 2). Dot plots show the percentage of nuclei in which a gene was detected and log(1 + mean normalised expression), scaled for each gene across clusters. Within the pioneer cluster, co-detection of NTN1, ITGAV and ITGB8 in the same nuclei was tested pairwise with Cochran-Mantel-Haenszel tests stratified by pool and by UMI-depth quintile within pool, with Benjamini-Hochberg adjustment across the three pairs. Bioinformatics pipeline references are in Supplemental Materials 1.

### Inducible HEK293 cell lines

To generate inducible NTN1 and eGFP stable cell lines with single site integrations, we first used Gateway cloning (Invitrogen) to construct FRT flanked expression vectors from human NTN1 with a C-terminal FLAG tag (#RC-218429; Origene) and a synthesised eGFP-FLAG control (Supplementary data). We prepared stable cells with integrated NTN1-FLAG or GFP-FLAG into the Flp-In^TM^ T-REx^TM^ parent cell line (R78007, Invitrogen) according to the manufacturer’s instructions, selecting clonal lines for all work presented. Cells were validated by Western blot or immunofluorescence against FLAG (Suppl.Fig.S3).

### Co-Immunoprecipitation

Stable HEK293 cells were transfected with plasmids encoding ITGB8-V5 and tetracycline (final concentration 1 µg/ml) was added 6 hours later to induce Netrin-1 expression. While A549 cells were transfected with NTN1(FLAG)-T2A-eGFP, ITGB8-V5, or FLAG-eGFP (plasmid maps and insert sequences are provided in Supplemental Materials 1). Transfections were performed in 6-well plates using 1 µg of total DNA per well and Lipofectamine LTX according to the manufacturer’s instructions. At 18 h post-transfection, cells had media removed and two gentle warm PBS washes and then lysed in ice cold NP-40 IP lysis buffer (150 mM NaCl, 1% NP-40, 50 mM Tris-HCl pH 8.0, 0.15% [w/v] BSA, 10% [v/v] glycerol, supplemented with Roche cOmplete EDTA-free protease inhibitor cocktail immediately prior to use). Samples in lysis buffer were then kept on ice with gentle agitation for 15 minutes before being scraped into a microfuge tube and clarified by centrifugation (12,000 xg for 30 minutes at 4 °C). Protein concentrations were determined by BCA microplate assay. With a small aliquot of each sample reserved as total input, equal amounts of lysate were incubated with end over end agitation for 4 hours at 4 °C and then subjected to coimmunoprecipitation using either Pierce Anti-DYKDDDDK Magnetic Agarose (A36797, Invitrogen) FLAG-tagged bait or Anti-V5 Magnetic Beads (SAE0203, Sigma) for v5-tagged bait. The manufacturers’ protocols were followed for both systems, except 50µl of beads was used; incubation with beads was at room temperature with end over end agitation for 1 hour; and final elution was performed using 1x NuPAGE LDS loading buffer and 1x NuPAGE reducing agent, at 95 °C for 3 minutes. Western blotting was then performed on the enriched samples alongside total input for comparison.

## DECLARATIONS

### Ethics approval and consent to participate

GE chickens were generated and bred under Home Office Project Licence PP1196147 with approval of the local AWERB committee. Chicken embryos at non-protected embryonic stages under the Animals (Scientific Procedures) Act 1986 were used for this study. Human data was not used in this study.

### Author Contributions

Conceptualization JR; methodology, TR, HH, TK, RQ, DD, MJM, DM, AR, JR; software, TK, RQ, DD; training and supervision, HH, AR, RQ, JR; investigation, TR, HH, HP, TK, RQ, DD, AJ, JI, MJM, DM, JR; formal analysis, TR, DD, HH and JR; resources, RQ, JR; data curation, TR, DD, RQ, TK; writing—original draft preparation, JR; writing—review and editing, TR, DD; funding acquisition, JR. All authors have read and agreed to the published version of the manuscript.

### Data and resource availability

Raw sequencing data are available in the ArrayExpress database (www.ebi.ac.uk/arrayexpress, brokered to the European Nucleotide Archive (ENA)), accession number E-MTAB-17975. All custom codes are available freely upon request.

## Acknowledgments

We would like to thank Samantha Bulter (UCLA) for helpful manuscript discussions, the staff at the Greenwood Building (National Avian Research Facility, Roslin Institute) for chicken husbandry, Anna Raper and colleagues at the Bioimaging and Flow Cytometry Facility (Roslin Institute) for cell sorting and imaging; and Steven Lisgo, Tamilvendhan Dhanaseelan, and team at the Newcastle University Genomics Core Facility and Bioinformatics Core for snRNAseq processing and initial analysis.

## Declaration of generative AI and AI-assisted technologies

During the preparation of this work, the author(s) used a large language model (Claude, Anthropic) to improve clarity and grammar. Bioinformatics methods were written with AI assistance (Claude, Anthropic), and were reviewed and checked by DD. The authors reviewed and edited the content as needed and take full responsibility for the content of the publication.

