## Supplemental Figures for "A gene-edited chicken model of ocular coloboma reveals Netrin-1 is required for pioneer cell detachment during optic fissure closure"

### SUPPLEMENTAL FIGURES & LEGENDS

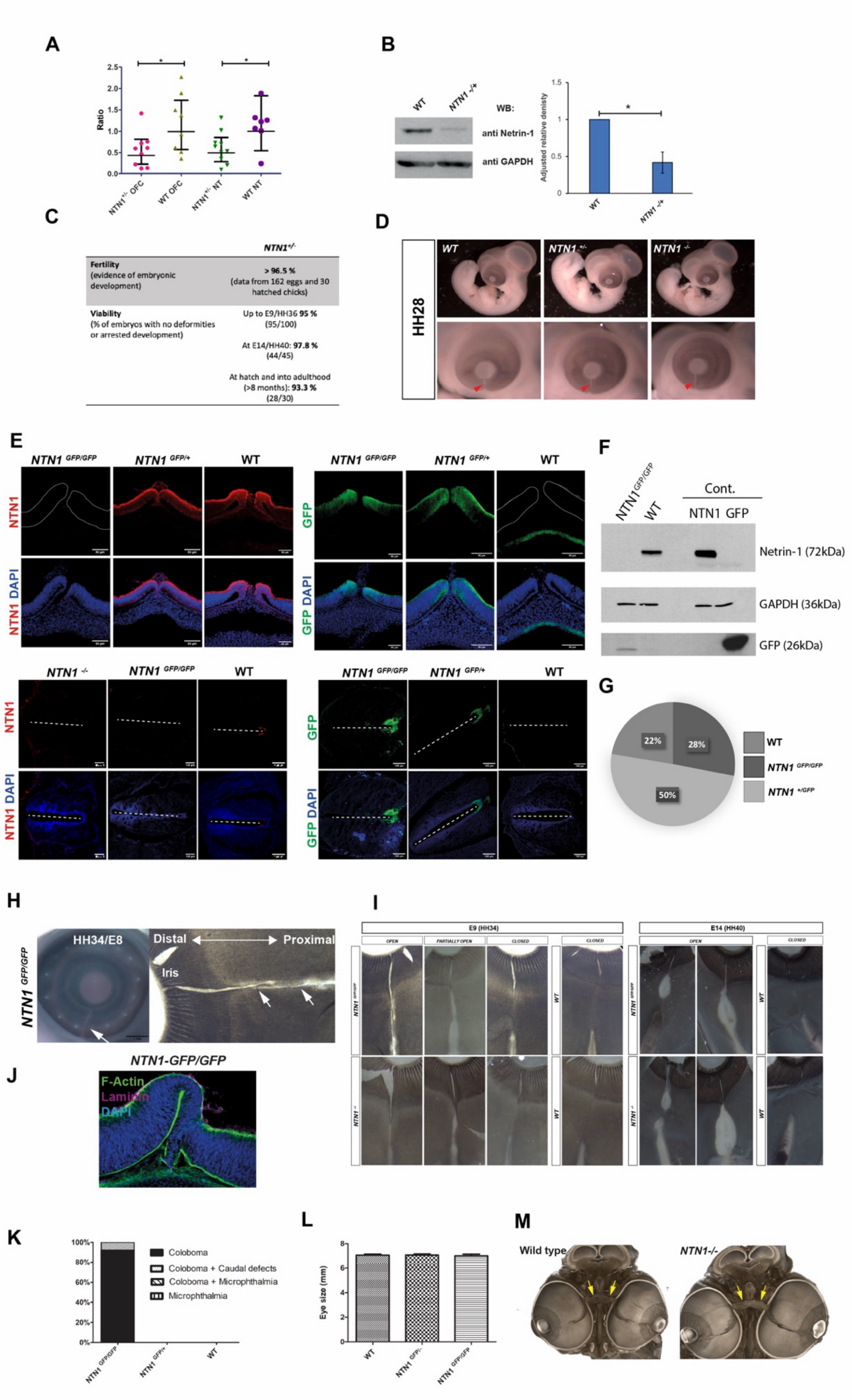

***Supplemental Figure S1.*** (**A**) Quantitative RT-PCR analysis of *NTN1* mRNA in tissues excised from OFC or neural tube (NT) at HH28. Both tissues show ~50 % reduction in transcript abundance for heterozygous $NTN1^{+/-}$ samples compared to WT control samples. (**B**) Protein analysis for Netrin-1 in optic fissures dissected at HH28 by western blot indicated ~50 % reduction in detectable protein in $NTN1^{+/-}$ compared to equivalent WT tissues. Quantitative data is representative of 3 independent dissections and analysis of 3 separate western blots. (**C**) Table showing fertility and viability of heterozygous $NTN1^{+/-}$ G1 embryos at multiple stages of development and hatching. (**D**) No deleterious phenotypes were observed in embryos collected at E6/HH28 across the three genotypes. (**E**) Immunofluorescence for Netrin-1 and GFP in $NTN1^{T2A\text{-}GFP/\,T2A\text{-}GFP}$ embryos compared to $NTN1^{T2A\text{-}GFP/+}$ and controls in the optic fissure and neural tube floorplate showed reciprocal staining and loss of detectable Netrin-1 in homozygotes. (**F**) Confirmation for loss of detectable Netrin-1 in $NTN1^{T2A\text{-}GFP/\,T2A\text{-}GFP}$ homozygotes by western blot from dissected optic fissure tissues. (**G**) Distribution of genotypes identified in heterozygote x heterozygote ($NTN1^{T2A\text{-}GFP/+}$x $NTN1^{T2A\text{-}GFP/+}$) crosses showed mendelian ratios for the three genotypes in collected embryos and hatched birds (*n* >250 birds genotyped). (**H**) Typical gross appearance of coloboma phenotype identified in $NTN1^{T2A\text{-}GFP/T2A\text{-}GFP}$ homozygotes. (**I**) Range of coloboma phenotypes observed in flat-mounted retinas from the two Netrin-1 deficient lines. (**J**) F-actin and Laminin immunofluorescence in $NTN1^{T2A\text{-}GFP/\,T2A\text{-}GFP}$ colobomas indicated these were phenotypically identical to $NTN1^{-/-}$ colobomas (n=3). (**K**) Penetrance of ocular phenotypes observed in *NTN1-GFP* line. (**L**) Size measurements of eye globes in homozygote embryos at E9 for both lines showed no difference to WT control eye sizes. (**M**) µCT analyses showed no gross structural differences of optic nerve and tract (arrows) in $NTN1^{-/-}$embryos (*n*=3 per genotype). * *P* <0.05, Student's t-test.

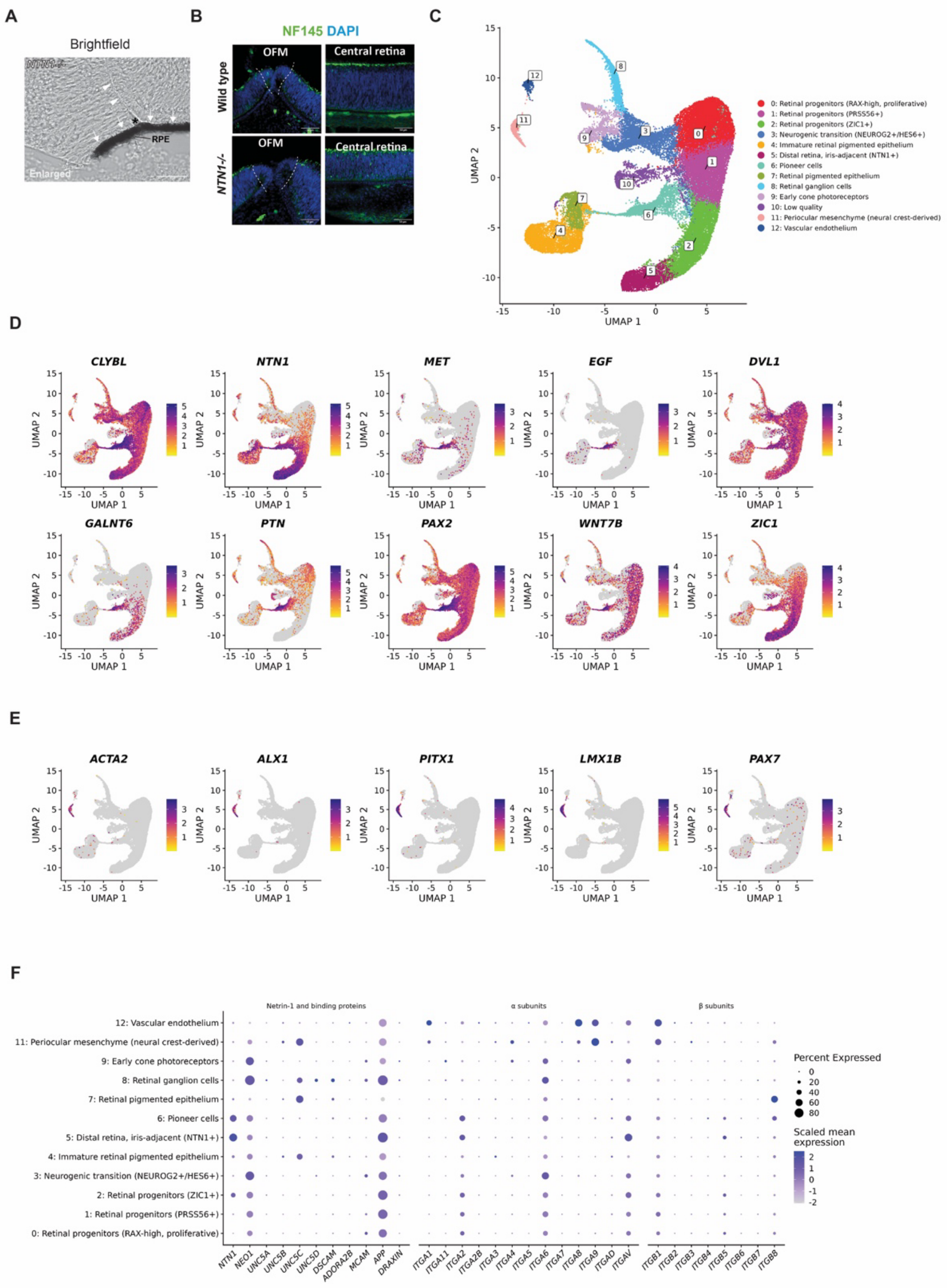


***Supplemental Figure S2.*** (**A**) Enlarged brightfield image of one side of a *NTN1*$^{-/-}$ fissure margin at HH40 indicating that RPE is a continuum (arrows) to which the overgrowth was attached, except for the internal aspect of the fold (arrowheads) which corresponds to a crease, where the asterisk indicates a folding point. No melanin granules were evident in the overgrown tissue. Scale bar = 25 µm. (**B**) Immunofluorescence for NF145 in mutant and WT at HH30 in pre-fusing fissures. No NF145 was observed in the pioneer cell region for either genotype. Scale bars = 50 µm. (**C**) UMAP of 46,930 nuclei from four pools of WT HH30 optic fissures, coloured by the 13 clusters. Clusters were labelled by comparing their marker genes with published pioneer cell markers (Trejo-Reveles et al., 2023; Chan et al., 2025) and literature markers of other ocular cell types. Cluster 6 was identified as pioneer cells and cluster 11 as periocular mesenchyme; cluster 10 (low quality) is shown in C to E but left out of F. (**D**, **E**) Expression of pioneer cell (**D**) and periocular mesenchyme (**E**) marker genes on the UMAP (log-normalised; grey, not detected; each gene on its own scale). (**F**) *NTN1*, its known binding proteins and every integrin subunit with a gene name in the Ensembl annotation of the chicken genome (GRCg7b), by cluster, cluster 10 excluded. Dot size: % of nuclei with the gene detected; colour: log(1 + mean normalised expression), scaled for each gene across clusters.

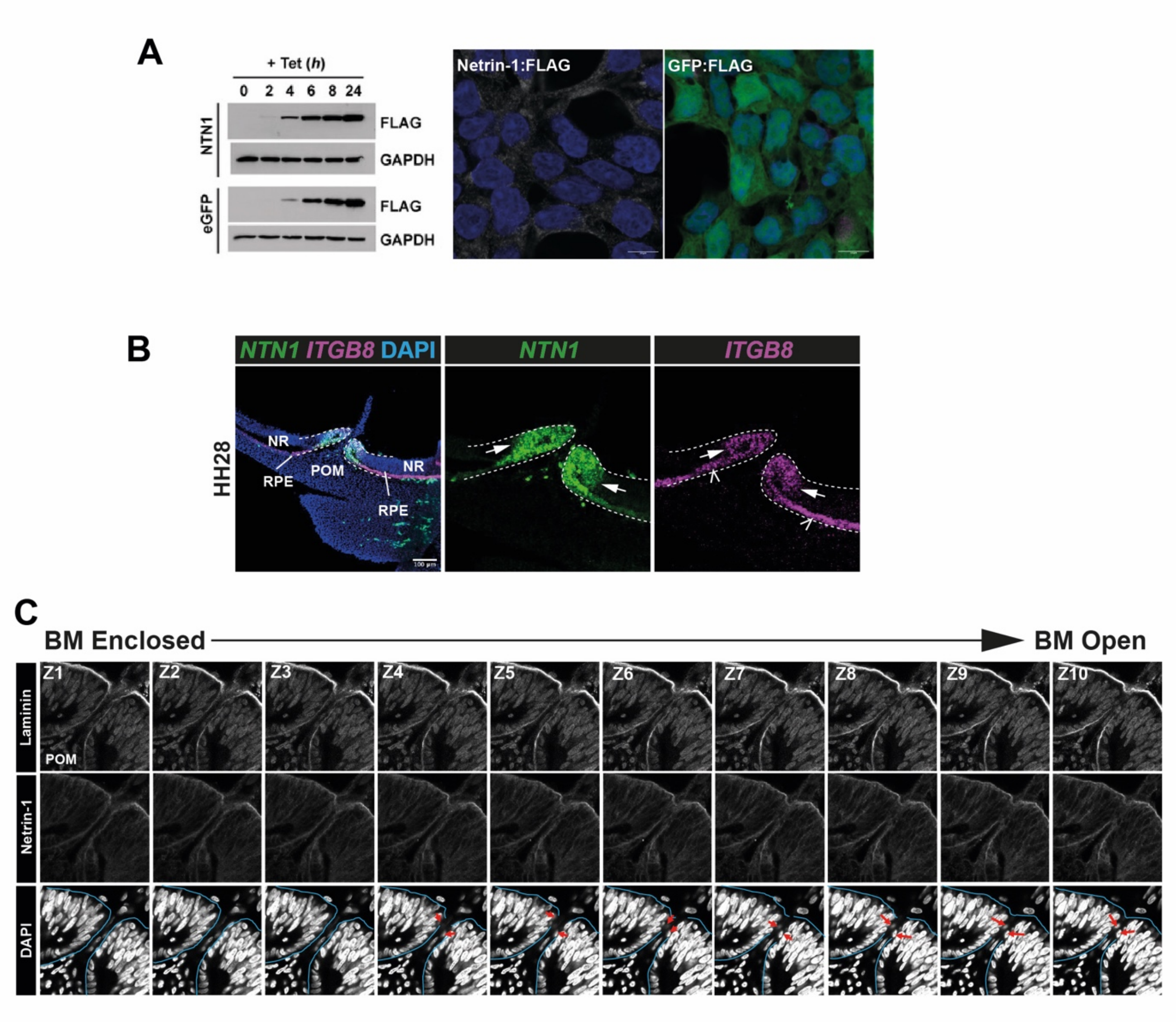


*Supplemental Figure S3.* (**A**) Western blot to validate inducible NTN1:FLAG and GFP:FLAG HEK293 Flp-In cell lines. Induced protein expression was detectable at 4 hours after tetracycline induction, consistent with their detection by immunofluorescence with anti-FLAG antibodies. (**B**) Fluorescence *in situ* hybridisation for *ITGB8* and *NTN1* in WT HH28/E6 optic fissures 24h prior to fusion. *NTN1* was present in pioneer cells (arrows); *ITGB8* was expressed in pioneer cells (arrows) and RPE (open arrowheads). (**C**) Consecutive confocal Z-stack slices for immunofluorescence against laminin, Fibronectin, and DAPI indicating mixing of fibronectin and laminin fragments adjacent to pioneer cells during fusion in WT fissures.
