## Supplemental Materials 1 for "A gene-edited chicken model of ocular coloboma reveals Netrin-1 is required for pioneer cell detachment during optic fissure closure"

**SUPPLEMENTAL INFORMATION –**

**Plasmids**

1. **Synthesised NTN1-T2A-eGFP**

pRP[Exp]-Puro-CAG>hNTN1[NM_004822.3](ns)/FLAG:T2A:EGFP

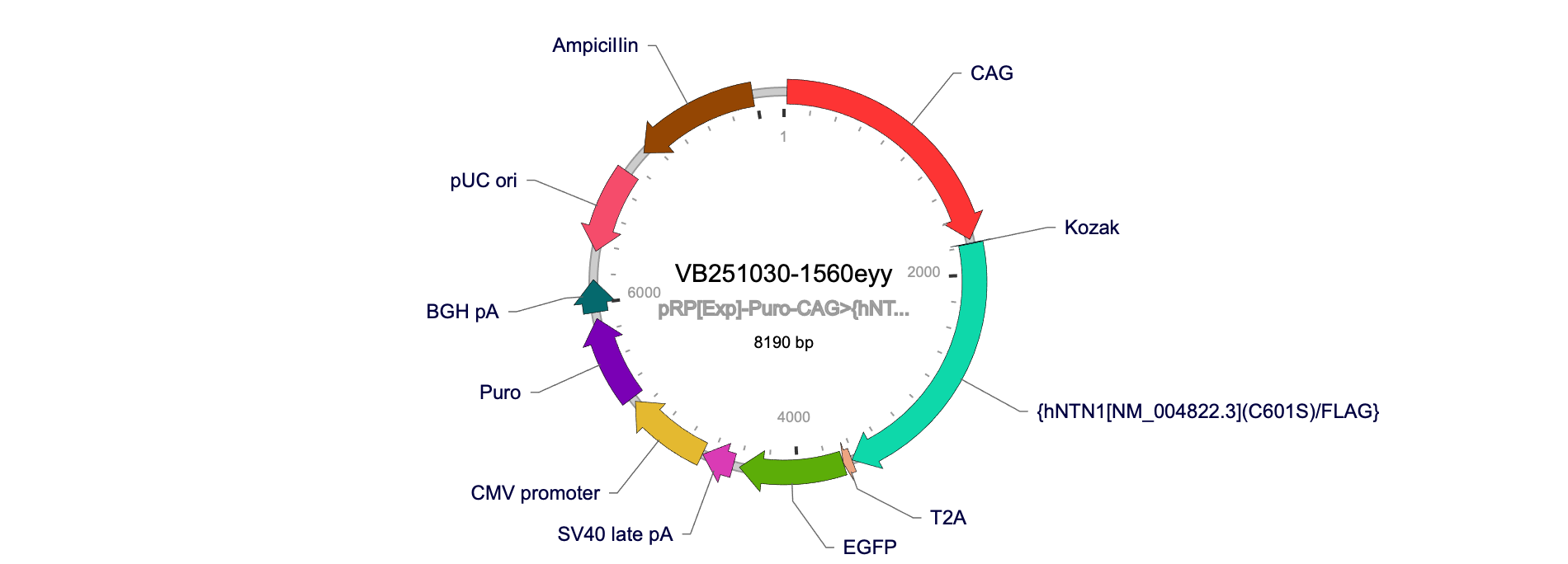

hNTN1[NM_004822.3](ns)/FLAG:T2A:EGFP

ATGATGCGCGCAGTGTGGGAGGCGCTGGCGGCGCTGGCGGCGGTGGCGTGCCTGGTGGGCGCGGTGCGCGGCGGGCCCGGGCTCAGCATGTTCGCGGGCCAGGCGGCGCAGCCCGATCCCTGCTCGGACGAGAACGGCCACCCGCGCCGCTGCATCCCGGACTTTGTCAATGCGGCCTTCGGCAAGGACGTGCGCGTGTCCAGCACCTGCGGCCGGCCCCCGGCGCGCTACTGCGTGGTGAGCGAGCGCGGCGAGGAGCGGCTGCGCTCGTGCCACCTCTGCAACGCGTCCGACCCCAAGAAGGCGCACCCGCCCGCCTTCCTCACCGACCTCAACAACCCGCACAACCTGACGTGCTGGCAGTCCGAGAACTACCTGCAGTTCCCGCACAACGTCACGCTCACACTGTCCCTCGGCAAGAAGTTCGAAGTGACCTACGTGAGCCTGCAGTTCTGCTCGCCGCGGCCCGAGTCCATGGCCATCTACAAGTCCATGGACTACGGGCGCACGTGGGTGCCCTTCCAGTTCTACTCCACGCAGTGCCGCAAGATGTACAACCGGCCGCACCGCGCGCCCATCACCAAGCAGAACGAGCAGGAGGCCGTGTGCACCGACTCGCACACCGACATGCGCCCGCTCTCGGGCGGCCTCATCGCCTTCAGCACGCTGGACGGGCGGCCCTCGGCGCACGACTTCGACAACTCGCCCGTGCTGCAGGACTGGGTCACGGCCACAGACATCCGCGTGGCCTTCAGCCGCCTGCACACGTTCGGCGACGAGAACGAGGACGACTCGGAGCTGGCGCGCGACTCGTACTTCTACGCGGTGTCCGACCTGCAGGTGGGCGGCCGGTGCAAGTGCAACGGCCACGCGGCCCGCTGCGTGCGCGACCGCGACGACAGCCTGGTGTGCGACTGCAGGCACAACACGGCCGGCCCGGAGTGCGACCGCTGCAAGCCCTTCCACTACGACCGGCCCTGGCAGCGCGCCACAGCCCGCGAAGCCAACGAGTGCGTGGCCTGTAACTGCAACCTGCATGCCCGGCGCTGCCGCTTCAACATGGAGCTCTACAAGCTTTCGGGGCGCAAGAGCGGAGGTGTCTGCCTCAACTGTCGCCACAACACCGCCGGCCGCCACTGCCATTACTGCAAGGAGGGCTACTACCGCGACATGGGCAAGCCCATCACCCACCGGAAGGCCTGCAAAGCCTGTGATTGCCACCCTGTGGGTGCTGCTGGCAAAACCTGCAACCAAACCACCGGCCAGTGTCCCTGCAAGGACGGCGTGACGGGTATCACCTGCAACCGCTGCGCCAAAGGCTACCAGCAGAGCCGCTCTCCCATCGCCCCCTGCATAAAGATCCCTGTAGCGCCGCCGACGACTGCAGCCAGCAGCGTGGAGGAGCCTGAAGACTGCGATTCCTACTGCAAGGCCTCCAAGGGGAAGCTGAAGATTAACATGAAAAAGTACTGCAAGAAGGACTATGCCGTCCAGATCCACATCCTGAAGGCGGACAAGGCGGGGGACTGGTGGAAGTTCACGGTGAACATCATCTCCGTGTATAAGCAGGGCACGAGCCGCATCCGCCGCGGTGACCAGAGCCTGTGGATCCGCTCGCGGGACATCGCCTGCAAGTGTCCCAAAATCAAGCCCCTCAAGAAGTACCTGCTGCTGGGCAACGCGGAGGACTCTCCGGACCAGAGCGGCATCGTGGCCGATAAAAGCAGCCTGGTGATCCAGTGGCGGGACACGTGGGCGCGGCGGCTGCGCAAGTTCCAGCAGCGTGAGAAGAAGGGCAAGTGCAAGAAGGCCGACTACAAAGACGATGACGACAAGTAG

2. **Synthesised eGFP-FLAG control**

pRP[Exp]-Puro-CAG>{EGFP-FLAG}

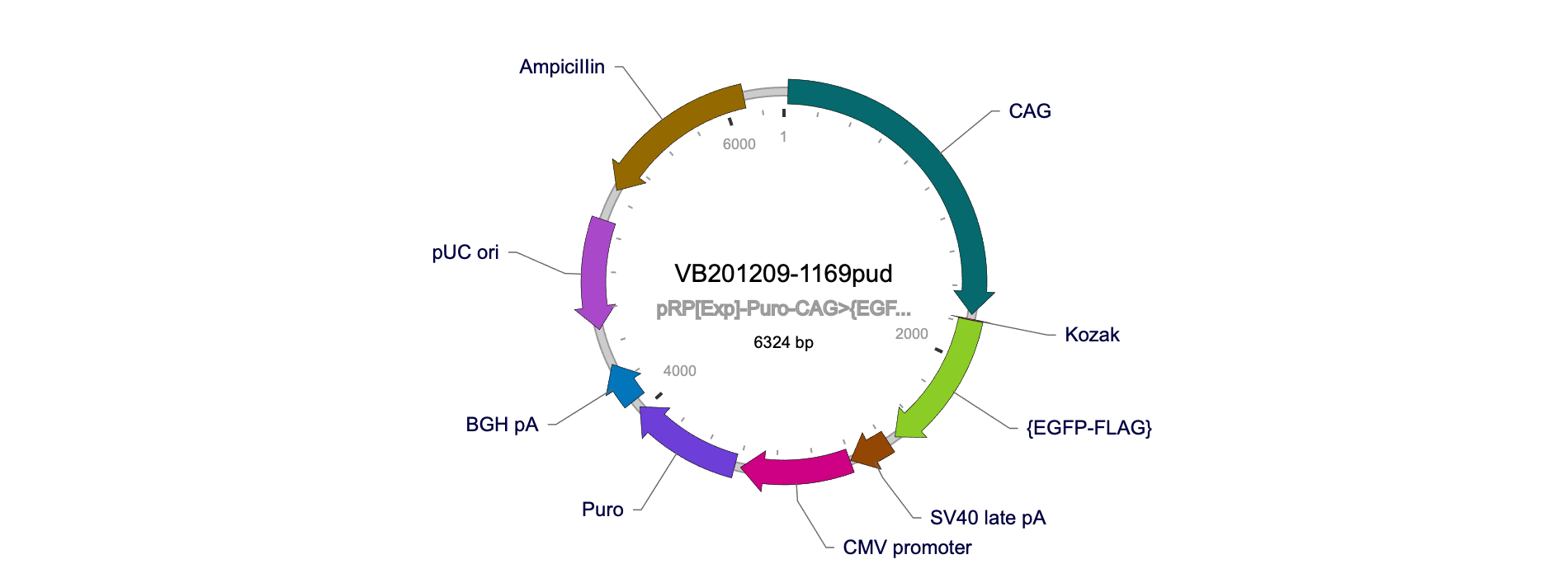

EGFP-FLAG

ATGGTGAGCAAGGGCGAGGAGCTGTTCACCGGGGTGGTGCCCATCCTGGTCGAGCTGGACGGCGACGTAAACGGCCACAAGTTCAGCGTGTCCGGCGAGGGCGAGGGCGATGCCACCTACGGCAAGCTGACCCTGAAGTTCATCTGCACCACCGGCAAGCTGCCCGTGCCCTGGCCCACCCTCGTGACCACCCTGACCTACGGCGTGCAGTGCTTCAGCCGCTACCCCGACCACATGAAGCAGCACGACTTCTTCAAGTCCGCCATGCCCGAAGGCTACGTCCAGGAGCGCACCATCTTCTTCAAGGACGACGGCAACTACAAGACCCGCGCCGAGGTGAAGTTCGAGGGCGACACCCTGGTGAACCGCATCGAGCTGAAGGGCATCGACTTCAAGGAGGACGGCAACATCCTGGGGCACAAGCTGGAGTACAACTACAACAGCCACAACGTCTATATCATGGCCGACAAGCAGAAGAACGGCATCAAGGTGAACTTCAAGATCCGCCACAACATCGAGGACGGCAGCGTGCAGCTCGCCGACCACTACCAGCAGAACACCCCCATCGGCGACGGCCCCGTGCTGCTGCCCGACAACCACTACCTGAGCACCCAGTCCGCCCTGAGCAAAGACCCCAACGAGAAGCGCGATCACATGGTCCTGCTGGAGTTCGTGACCGCCGCCGGGATCACTCTCGGCATGGACGAGCTGTACAAGGGAAGCGGAGACTACAAAGACGATGACGACAAGTAA

3. **Synthesised ITGB8-V5**

pRP[Exp]-Puro-CAG>hITGB8[NM_002214.3](ns)*/3xGGGGS/V5

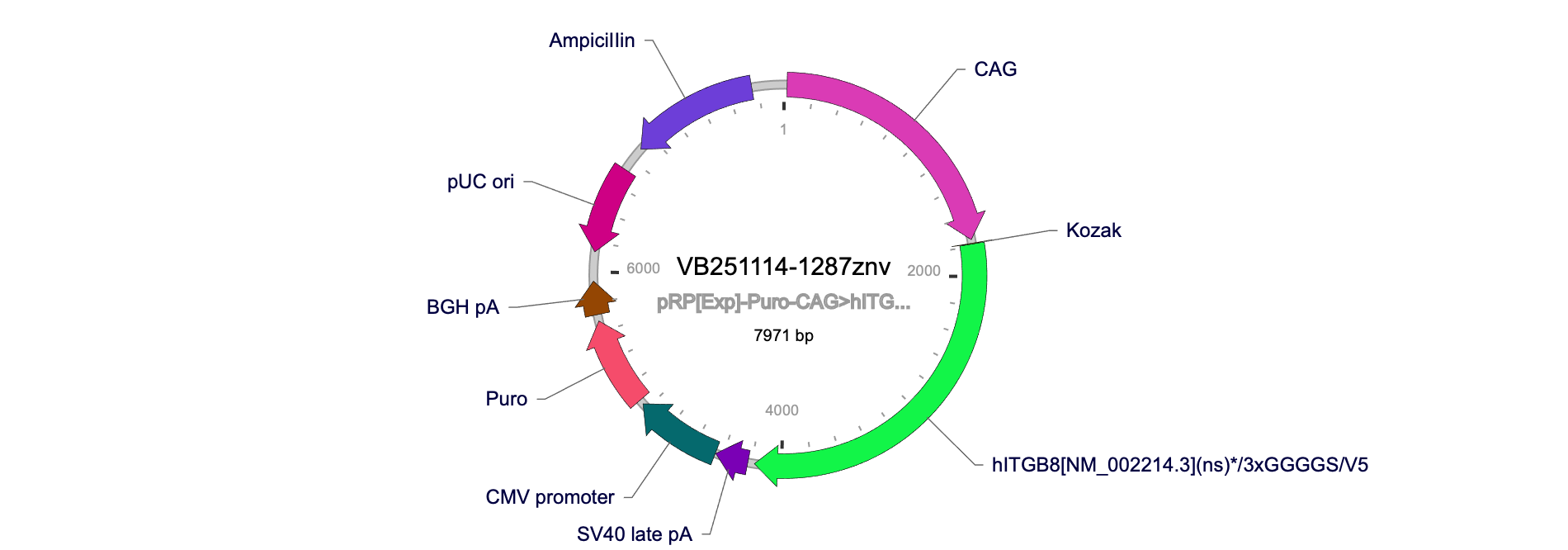

hITGB8[NM_002214.3](ns)*/3xGGGGS/V5

ATGTGCGGCTCGGCCCTGGCTTTTTTTACCGCTGCATTTGTCTGCCTGCAAAACGACCGGCGAGGTCCCGCCTCGTTCCTCTGGGCAGCCTGGGTGTTTTCACTTGTTCTTGGACTGGGCCAAGGTGAAGACAATAGATGTGCATCTTCAAATGCAGCATCCTGTGCCAGGTGCCTTGCGCTGGGTCCAGAATGTGGATGGTGTGTTCAAGAGGATTTCATTTCAGGTGGATCAAGAAGTGAACGTTGTGATATTGTTTCCAATTTAATAAGCAAAGGCTGCTCAGTTGATTCAATAGAATACCCATCTGTGCATGTTATAATACCCACTGAAAATGAAATTAATACCCAGGTGACACCAGGAGAAGTGTCTATCCAGCTGCGTCCAGGAGCCGAAGCTAATTTTATGCTGAAAGTTCATCCTCTGAAGAAATATCCTGTGGATCTTTATTATCTTGTTGATGTCTCAGCATCAATGCACAATAATATAGAAAAATTAAATTCCGTTGGAAACGATTTATCTAGAAAAATGGCATTTTTCTCCCGTGACTTTCGTCTTGGATTTGGCTCATACGTTGATAAAACAGTTTCACCATACATTAGCATCCACCCCGAAAGGATTCATAATCAATGCAGTGACTACAATTTAGACTGCATGCCTCCCCATGGATACATCCATGTGCTGTCTTTGACAGAGAACATCACTGAGTTTGAGAAAGCAGTTCATAGACAGAAGATCTCTGGAAACATAGATACACCAGAAGGAGGTTTTGACGCCATGCTTCAGGCAGCTGTCTGTGAAAGTCATATCGGATGGCGAAAAGAGGCTAAAAGATTGCTGCTGGTGATGACAGATCAGACGTCTCATCTCGCTCTTGATAGCAAATTGGCAGGCATAGTGGTGCCCAATGACGGAAACTGTCATCTGAAAAACAACGTCTATGTCAAATCGACAACCATGGAACACCCCTCACTAGGCCAACTTTCAGAGAAATTAATAGACAACAACATTAATGTCATCTTTGCAGTTCAAGGAAAACAATTTCATTGGTATAAGGATCTTCTACCCCTCTTGCCAGGCACCATTGCTGGTGAAATAGAATCAAAGGCTGCAAACCTCAATAATTTGGTAGTGGAAGCCTATCAGAAGCTCATTTCAGAAGTGAAAGTTCAGGTGGAAAACCAGGTACAAGGCATCTATTTTAACATTACCGCCATCTGTCCAGATGGGTCCAGAAAGCCAGGCATGGAAGGATGCAGAAACGTGACGAGCAATGATGAAGTTCTTTTCAATGTAACAGTTACAATGAAAAAATGTGATGTCACAGGAGGAAAAAACTATGCAATAATCAAACCTATTGGTTTTAATGAAACCGCTAAAATTCATATACACAGAAACTGCAGCTGTCAGTGTGAGGACAACAGAGGACCTAAAGGAAAGTGTGTAGATGAAACTTTTCTAGATTCCAAGTGTTTCCAGTGTGATGAGAATAAATGTCATTTTGATGAAGATCAGTTTTCTTCTGAGAGTTGCAAGTCACACAAGGATCAGCCTGTTTGCAGTGGTCGAGGAGTTTGTGTTTGTGGGAAATGTTCATGTCACAAAATTAAGCTTGGAAAAGTGTATGGAAAATACTGTGAAAAGGATGACTTTTCTTGTCCATATCACCATGGAAATCTGTGTGCTGGGCATGGAGAGTGTGAAGCAGGCAGATGCCAATGCTTCAGTGGCTGGGAAGGTGATCGATGCCAGTGCCCTTCAGCAGCAGCCCAGCACTGTGTCAATTCAAAGGGCCAAGTGTGCAGTGGAAGAGGCACGTGTGTGTGTGGAAGGTGTGAGTGCACCGATCCCAGGAGCATCGGCCGCTTCTGTGAACACTGCCCCACCTGTTATACAGCCTGCAAGGAAAACTGGAATTGTATGCAATGCCTTCACCCTCACAATTTGTCTCAGGCTATACTTGATCAGTGCAAAACCTCATGTGCTCTCATGGAACAACAGCATTATGTCGACCAAACTTCAGAATGTTTCTCCAGCCCAAGCTACTTGAGAATATTTTTCATCATTTTCATAGTTACATTCTTGATTGGGTTGCTTAAAGTCCTGATCATTAGACAGGTGATACTACAATGGAATAGTAATAAAATTAAGTCCTCATCAGATTACAGAGTGTCAGCCTCAAAAAAGGATAAGTTGATTCTGCAAAGTGTTTGCACAAGAGCAGTCACCTACCGACGTGAGAAGCCTGAAGAAATAAAAATGGATATCAGCAAATTAAATGCTCATGAAACTTTCAGGTGCAACTTCTCTGGTGGCGGAGGCTCGGGCGGAGGTGGGTCGGGTGGCGGCGGATCAGGTAAGCCTATCCCTAACCCTCTCCTCGGTCTCGATTCTACGTAA

**SUPPLEMENTAL INFORMATION -**

**snRNAseq bioinformatics pipeline references**

Ahlmann-Eltze C, Huber W. glmGamPoi: fitting Gamma-Poisson generalized linear models on single cell count data. Bioinformatics. 2021;36:5701-5702.

Amezquita RA, Lun ATL, Becht E, et al. Orchestrating single-cell analysis with Bioconductor. Nat Methods. 2020;17:137-145.

Amezquita RA, Lun ATL, Hicks SC, Gottardo R. Orchestrating Single-Cell Analysis with Bioconductor (online book, Bioconductor 3.23 release). 2026. https://bioconductor.org/books/release/OSCA/

Choudhary S, Satija R. Comparison and evaluation of statistical error models for scRNA-seq. Genome Biol. 2022;23:27.

Ewels PA, Peltzer A, Fillinger S, et al. The nf-core framework for community-curated bioinformatics pipelines. Nat Biotechnol. 2020;38:276-278.

Friocourt F, Lafont AG, Kress C, et al. Recurrent DCC gene losses during bird evolution. Sci Rep. 2017;7:37569.

Fleming SJ, Chaffin MD, Arduini A, et al. Unsupervised removal of systematic background noise from droplet-based single-cell experiments using CellBender. Nat Methods. 2023;20:1323-1335.

Germain PL, Lun A, Garcia Meixide C, Macnair W, Robinson MD. Doublet identification in single-cell sequencing data using scDblFinder. F1000Res. 2021;10:979.

Hafemeister C, Satija R. Normalization and variance stabilization of single-cell RNA-seq data using regularized negative binomial regression. Genome Biol. 2019;20:296.

Hao Y, Stuart T, Kowalski MH, et al. Dictionary learning for integrative, multimodal and scalable single-cell analysis. Nat Biotechnol. 2024;42:293-304.

Korsunsky I, Millard N, Fan J, et al. Fast, sensitive and accurate integration of single-cell data with Harmony. Nat Methods. 2019;16:1289-1296.

Love MI, Huber W, Anders S. Moderated estimation of fold change and dispersion for RNA-seq data with DESeq2. Genome Biol. 2014;15:550.

Tirosh I, Izar B, Prakadan SM, et al. Dissecting the multicellular ecosystem of metastatic melanoma by single-cell RNA-seq. Science. 2016;352:189-196.

Zheng GXY, Terry JM, Belgrader P, et al. Massively parallel digital transcriptional profiling of single cells. Nat Commun. 2017;8:14049.
